# Pharmacological MCL-1 inhibition disrupts fatty acid oxidation and depletes neural progenitor cells

**DOI:** 10.64898/2025.12.12.694056

**Authors:** Marina R. Hanna, Madison Yarbrough, James Costanzo, Melanie Gil, Matthew Murrow, Vivian Gama

## Abstract

MCL-1 is a canonical anti-apoptotic protein crucial for early neurodevelopment, and its loss causes embryonic-lethal defects that other BCL-2 family proteins cannot rescue. Here, we pharmacologically inhibit MCL-1 in human neural progenitor cells and uncover non-apoptotic roles in sustaining mitochondrial cristae integrity, fatty acid oxidation, and progenitor identity. MCL-1 inhibition disrupts mitochondrial ultrastructure, destabilizing the OPA1-MICOS machinery. These structural defects are accompanied by ACSL1 displacement from the mitochondria, impaired fatty acid oxidation, lipid droplet accumulation, and reduced oxygen consumption, revealing a tight link between cristae architecture and metabolic competence. Mechanistically, MCL-1 acts through two coordinated functions: maintaining cristae integrity at the inner membrane and retaining ACSL1 at the outer membrane, both independently of caspase activation. Functionally, MCL-1 inhibition selectively depletes intermediate progenitor cells without affecting proliferation, indicating a direct role in lineage progression. Together, our findings position MCL-1 upstream of OPA1, MICOS, and ACSL1 as a critical coordinator of cristae organization, lipid metabolism, and neural progenitor fate, establishing mitochondrial inner membrane architecture as an instructive determinant of human neurogenesis and highlighting non-canonical MCL-1 functions as regulators of brain development.

## Introduction

Neurodevelopment relies on the precise coordination of cellular processes that determine the fate of multipotent *PAX6*-expressing radial glial cells (RGCs)^1–3^. RGCs are the major neural stem cells (NSCs) that give rise to glia and neurons in the human cortex^4,5^. The fate of these NSCs relies on their ability to undergo symmetric or asymmetric division, a process that ultimately determines whether they self-renew or differentiate into specialized cell types^4,5^. NSCs give rise to intermediate progenitor cells (IPCs)^6,7^, which are *EOMES* (TBR2)- expressing^8,9^ cortex-specific transit amplifying cells^10^ committed to producing glutamatergic projection neurons of cortical layers two through six^11,12^. Given the high bioenergetic demands and metabolic complexity of NSCs, disruption of mitochondrial morphology and function can impair critical stem cell transitions during neurogenesis^13–16^. Despite this, the precise role of mitochondrial morphology and function in these intricate transition states is not fully understood.

Myeloid cell leukemia-1 (MCL-1) is a mitochondrial membrane protein identified as a regulator of mitochondrial morphology through mechanisms that are independent of its canonical anti-apoptotic function^17–21^. Initially recognized only as a member of the B-cell lymphoma 2 (BCL- 2) family for its <u>B</u>CL-2 <u>h</u>omology-<u>3</u> (BH3) domain and role in preventing cell death at the outer mitochondrial membrane (OMM)^22^, MCL-1 has been well-established to have non-apoptotic functions^18,19,23–27^. Disruption of MCL-1 causes mitochondrial dysfunction in human cardiomyocytes^21,24^, as well as improper development of hematopoietic stem cells^28^, B cell and T cell progenitors^29^, activated B cells^30^, and the central nervous system^31^. Inhibition of MCL-1 has been explored as a promising treatment for chemo-resistant tumors but has not advance beyond Phase I/II in clinical trials due to potential cardiotoxicity. Inhibition of MCL-1 with existing small molecule inhibitors results in impaired cardiomyocyte function and disruption of mitochondrial morphology^32^, but more investigation is needed to determine whether this applies to other bioenergetically demanding cell types, such as, NSCs.

Previous work has reported a close association between MCL-1, mitochondrial morphology, and pluripotent stem cell identity of human embryonic stem cells (hESCs)^18^. Knockdown of *MCL-1* (*MCL-1^KD^*) in hESCs^18^ results in an elongated mitochondrial network, a phenotype linked to differentiation^33^. Moreover, *MCL-1^KD^* hESCs exhibited loss of key pluripotent transcription factors and identity markers, including *OCT4* and *NANOG*^34,35^. Other studies have established a postmitotic period of fate plasticity, controlled by mitochondrial morphology, during mouse brain development. When mitochondria remain elongated (low fission), NSCs divide symmetrically, thus expanding the pool of progenitor cells. Conversely, increased mitochondrial fission drives NSCs toward differentiation. *In vivo* mouse studies highlight the importance of MCL-1 in mouse embryonic survival and proper fetal neurodevelopment^27^. A gene swap approach in which the coding sequence of the *Mcl-1* gene was replaced with other anti-apoptotic genes, *Bcl-2* (*Mcl-1^Bcl-2^*), *Bcl-xL* (*Mcl-1^Bcl-xL^*), or *A1* (*Mcl-1^A1^*), resulted in neurodevelopmental defects that could not be rescued by anti-apoptotic functions alone, providing evidence that the noncanonical functions of MCL-1 are indispensable for proper neurodevelopment^27^.

MCL-1 has also been found to localize to the inner mitochondrial membrane (IMM), where it may interact with cristae-shaping proteins, such as optic atrophy type 1 (OPA1)^19,20,24^. However, the function of MCL-1 at the IMM remains poorly delineated. Studies in the field have shown that disruption of the fatty acid oxidation (FAO) pathway promotes asymmetric differentiation of adult mouse NSCs, highlighting the role of FAO in regulating NSC fate decisions^36–39^. Related work in cancer cells from the Opferman and Walensky groups revealed an unexpected link between MCL-1 and FAO^23,25,26^. Inhibition of MCL-1 results in decreased oxidation of long-chain fatty acids (LCFAs) in cancer cells^23,25^, likely due to the disrupted interaction between MCL-1 and the BH3-like domain of acyl-CoA synthetase long-chain family member 1 (ACSL1). This suggests that MCL-1 is a direct modulator of FAO in cancer cells.

FAO is a metabolic process critical for energy production and cellular homeostasis that occurs within the mitochondria^40^. This pathway requires LCFAs to translocate from the cytosol and pass through the OMM and IMM to reach their destination for beta oxidation in the mitochondrial matrix. Starting at the OMM, an LCFA is first activated to acyl-CoA derivatives by ACSL1 and then catalyzed by carnitine palmitoyltransferase I (CPT1) into fatty acylcarnitine, which diffuses through the OMM and enters the intermembrane space. The fatty acylcarnitine is then shuttled through the IMM by carnitine-acylcarnitine translocase (CACT). Once at the matrix, carnitine palmitoyltransferase II (CPT2) cleaves the fatty acylcarnitine, catalyzing the LCFA back to fatty acyl-CoA for β-oxidation at the matrix initiated by Very Long Acyl-CoA Dehydrogenase (VLCAD). The enzymatic breakdown of LCFAs is a multistep process that produces acetyl-CoA, a metabolic byproduct which enters the tricarboxylic acid (TCA) cycle. Gene transcription is influenced by mitochondrial metabolism through several metabolic byproducts, such, acetyl-CoA, alpha-ketoglutarate, and succinate^41,42^. Disruption of the FAO pathway promotes asymmetric differentiation of pluripotent and neural stem cells^36–39,43^, suggesting a role of FAO in stem cell fate decisions. Although FAO modulates neurodevelopment and MCL-1 is known to impact FAO of cancer cells, the role of MCL-1 in FAO during human neurodevelopment has not been explored. Here, we report the non-apoptotic effects of MCL-1 inhibition on mitochondrial morphology, cristae architecture, and FAO in an *in vitro* model of early human neurogenesis. Our findings provide evidence that MCL-1 maintains the integrity of mitochondrial cristae and neural progenitor cell identity.

## Results

### MCL-1 inhibition induces mitochondrial remodeling and inner membrane re- organization in human neural progenitor cells

MCL-1 is known for its canonical function as an anti-apoptotic member of the BCL-2 family and is indispensable for the survival of human induced pluripotent stem cells^18^, hematopoietic stem cells^28,44^, and cardiomyocytes^24^. To examine the effect of MCL-1 inhibition on human neural progenitor cells (NPCs), we differentiated human embryonic stem cells (hESCs) using dual SMAD inhibition, as previously described^45^. Human NPCs were treated for 24 and 48 hours with S63845, a selective BH3 mimetic inhibitor of MCL-1, for 24 or 48 hours (**Figure 1A**). Cells were co-treated with Q-VD-OPh (QVD), a potent pan-caspase inhibitor, to distinguish between phenotypes that are attributed to MCL-1’s anti-apoptotic or non-apoptotic functions, and to capture the transient state of cell types that are more vulnerable to MCL-1 inhibition. Using real-time quantitative polymerase chain reaction (RT-qPCR), we detected downregulation of *MCL-1* with S63485 treatment at 24 hours but not 48 hours, while co-treatment with QVD upregulated *MCL-1* expression at both timepoints (**Supplementary Figure 1A**). Previous studies reported an increase in MCL-1 protein levels upon MCL-1 inhibition, which is speculated to result from stabilization of MCL-1, whereby it is protected from proteasomal degradation when its BH3 domain is occupied^46,47^. We probed for MCL-1 protein levels and confirmed an increase at both timepoints with S63845 treatment (**Supplementary Figure 1B**-**C**) despite the downregulation of *MCL-1* expression at 24 hours. This increase in MCL-1 protein levels serves as an internal control that the S63845 small molecule is effectively inhibiting MCL-1 in NPCs. Increased MCL-1 levels were not detected with QVD co-treatment despite upregulation of *MCL-1* with co-treatment at both timepoints (**Supplementary Figure 1A-C**), confirming that QVD co-treatment captures the state of a subset of NPCs that are more vulnerable to MCL-1 inhibition.

**Figure 1.**
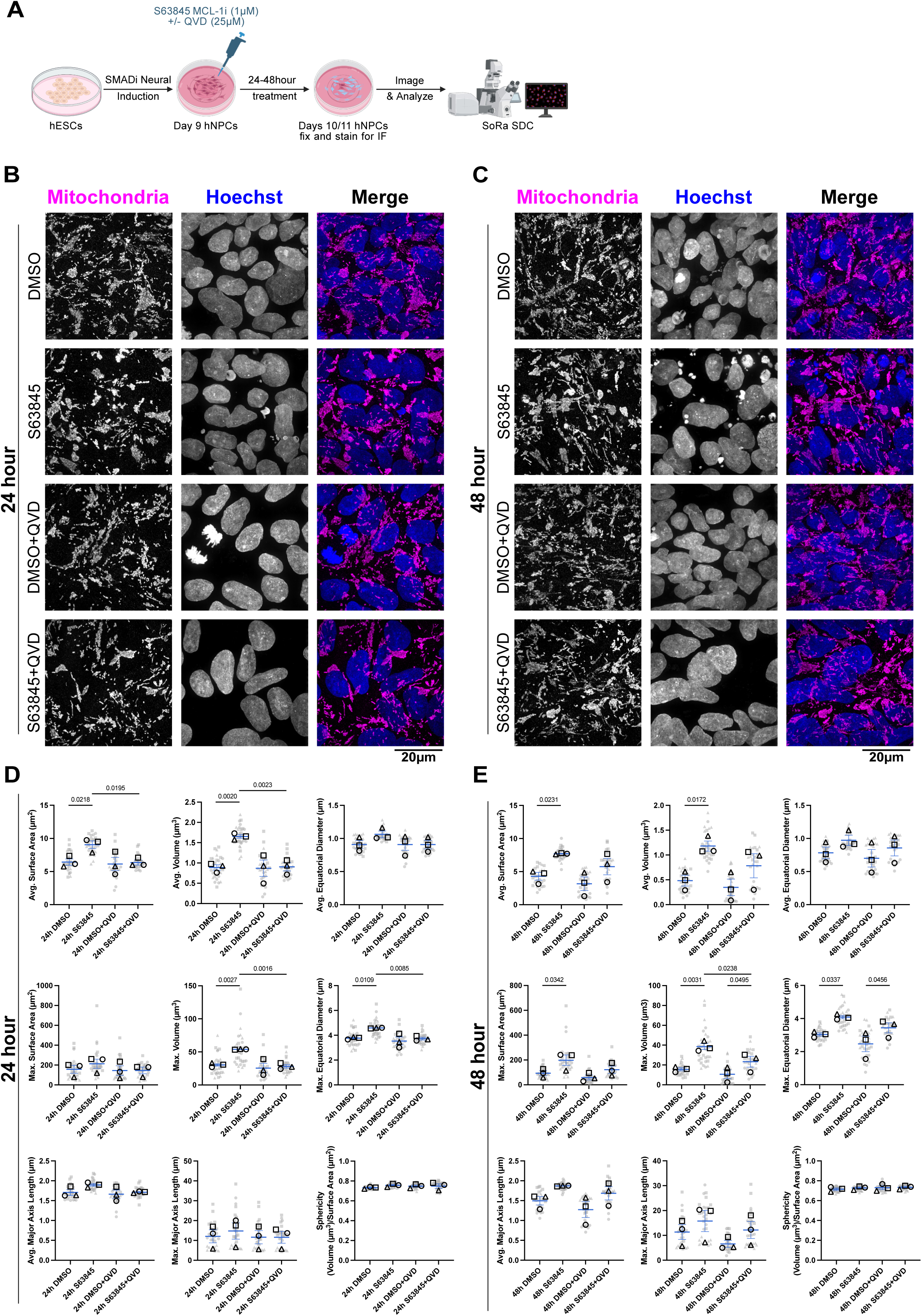
S63845 impacts mitochondrial content and morphology. (**A**) Schematic detailing the neural induction of human embryonic stem cells (hESCs) using SMAD inhibition (SMADi) for the derivation of the human neural progenitor cells (NPCs) tissue culture model. NPCs were maintained in neural induction media (NIM) with daily media changes until they were passaged on day 7. NPCs were given a 24-hour acclimation period on day 8 before treatment on day 9 with 1µm S63845 (MCL-1 inhibitor) −/+ 25µM QVD (caspase inhibitor). Cells were then fixed in paraformaldehyde at 24- and 48-hours post-treatment on days 10-11, respectively, and processed for imaging on a SoRa SDC (Super Resolution by Optical Reassignment Spinning Disk Confocal) microscope. (**B, C**) Immunofluorescent images of mitochondria (anti-mito in magenta) and nuclei (Hoechst in blue) acquired on SoRa SDC at 100X with 2.8X magnification (scale bar = 20µm). Representative images (n=3) of NPCs treated with S63845 (MCL-1 inhibitor) −/+ QVD (caspase inhibitor) for 24- (B) and 48-hours (**C**). (**D, E**) Quantified mitochondrial morphology parameters including sphericity, average and maximum surface area, volume, equatorial diameter, and major axis length for 24h (**D**) and 48h (**E**) treatments. Each large, opened-shape data point (black) represents the mean of a biological replicate, and each small, closed-shape data point (transparent gray) represents a technical replicate (a single image). Shape of the data points corresponds to a biological replicate (circle for n=1, square for n=2, and triangle for n=3). Data were analyzed using an ordinary two-way ANOVA with an uncorrected Fisher’s LSD test for multiple comparisons and a 95% confidence level (p-value ≤ 0.05 as the significance threshold). Error bars represent standard error of the mean.

MCL-1 inhibition is known to alter mitochondrial morphology in various cell types^18,32,49^. Thus, we first investigated the effects of S63845 on mitochondrial morphology in human NPCs. Super-resolution optical pixel reassignment (SoRa) microscopy revealed a pronounced swelling of the mitochondrial network in NPCs treated with S63845 at both timepoints (**Figure 1B-C**), cell viability was not overtly compromised by S63845 treatment, as evidenced by comparable nuclear morphology across all conditions and timepoints examined (**Figure 1B- C**). Mitochondrial morphology was quantified as previously described^15,32^. Consistent with these qualitative observations, average mitochondrial surface area and volume were significantly increased at both timepoints (**Figure 1D-E**). Moreover, maximum mitochondrial volume and equatorial diameter were significantly increased at both timepoints, indicating a globally expanded mitochondrial network. At 48 hours, this expansion was further reflected by a significant increase in maximum surface area in S63845-treated cells relative to controls (**Figure 1E**). Notably, this mitochondrial network expansion, rather than the fragmentation typically associated with apoptotic cell death, is consistent with a non-apoptotic cellular response to MCL-1 inhibition. To evaluate the contribution of caspase activity to these morphological changes, cells were co-treated with QVD. At 24 hours, co-treatment with QVD significantly abrogated the mitochondrial network expansion induced by S63845, with no significant differences observed relative to QVD-only controls (**Figure 1D**). In contrast, at 48 hours, the S63845-induced mitochondrial remodeling was maintained despite caspase inhibition, suggesting that the dependency on caspase activity for mitochondrial network expansion is temporally restricted to earlier stages of the cellular response (**Figure 1E**).

Given the critical role of mitochondrial cristae in metabolic regulation, we next investigated whether MCL-1 inhibition-induced alterations in mitochondrial morphology at 48 hours of S63845 treatment extended to cristae ultrastructure using transmission electron microscopy (TEM) and focused ion beam-scanning electron microscopy (FIB-SEM) (**Figure 2A**). TEM images revealed atypical cristae morphology in NPCs treated with S63845 for 48 hours, characterized by swollen matrices and sparse, disorganized cristae membranes, in contrast to the densely folded inner membrane architecture observed in control mitochondria (**Figure 2B**). To further characterize these ultrastructural changes in three dimensions, we performed volumetric reconstruction of mitochondrial cristae with FIB-SEM acquired images (**Figure 2C- E**). Three-dimensional analyses revealed significant increases in cristae surface area and volume in S63845-treated cells relative to controls (**Figure 2D**). Volumetric reconstruction of mitochondria (**Supplementary Figure 2A**) revealed that neither the mitochondrial branching index (MBI) nor sphericity were significantly affected (**Supplementary Figure 2B**). Collectively, these findings demonstrate that MCL-1 inhibition induces extensive disruption of cristae organization (**Figure 2A-E; Supplementary Videos 1 and 2**). These data implicate MCL-1 as a critical regulator of the structural integrity of the mitochondrial inner membrane.

**Figure 2.**
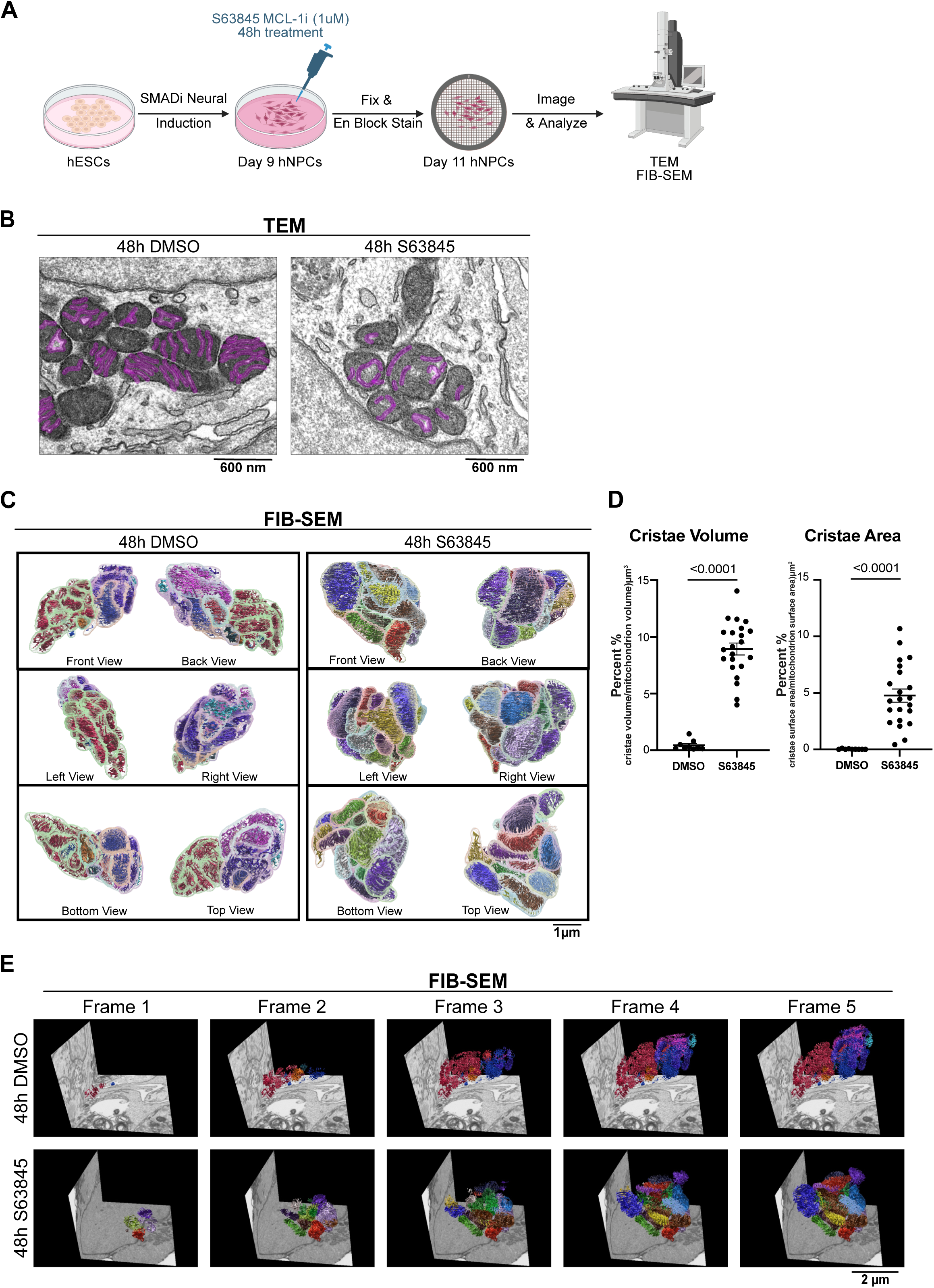
S63845 compromises mitochondrial cristae integrity. (**A**) Schematic detailing the neural induction of human embryonic stem cells (hESCs) using SMAD inhibition (SMADi) for the derivation of the human neural progenitor cells (NPCs) tissue culture model. NPCs were maintained in neural induction media (NIM) with daily media changes until they are passaged on day 7. NPCs were given a 24-hour acclimation period on day 8 before treatment on day 9 with 1µm S63845 (MCL-1 inhibitor). Cells were then collected at 48-hours post treatment on day 11 and processed for transmission electron microscopy (TEM) or focused ion beam-scanning electron microscopy (FIB-SEM). (**B**) Representative TEM images (n=3) of NPC mitochondrial cristae 48-hours post treatment with DMSO control and S63845 MCL-1 inhibitor (scale bar = 600nm). (**C**) 3-dimensional reconstructions of mitochondria and cristae (scale bar = 0.5µm). (**D**) Quantification of cristae changes from 3-dimensional reconstruction. Each cristae volume and surface area was normalized to its corresponding mitochondrion volume and surface area, respectively, and multiplied by 100 to calculate the percentage of cristae-occupied mitochondrial volume or area. Each data point is an individual mitochondrion. Data were analyzed using a parametric two-tailed unpaired t-test with a Welch’s correction. (**E**) Volumetric still frames of mitochondrial cristae 3-dimensional reconstruction of NPCs at 48 hours post treatment with DMSO control and S63845 (MCL-1 inhibitor) segmented from FIB-SEM volume images (scale bar = 2µm).

### MCL-1 activity preserves OPA1 proteostasis, MICOS complex stability, and mitochondrial respiration

Given the crista defects observed upon MCL-1 inhibition, we next examined key molecular regulators of cristae architecture and junction formation (**Figure 3A**). Cristae structure is maintained by the mitochondrial contact site and cristae-organizing system (MICOS), which comprises two core subcomplexes: the MIC60 subcomplex (MIC60, MIC19, MIC25), and the MIC10 subcomplex (MIC10, MIC26, MIC27), bridged by MIC13/QIL1^50^. At the inner mitochondrial membrane (IMM), MICOS establishes cristae junctions, which are required for cristae biogenesis and expansion of IMM surface area to support ATP synthesis via oxidative phosphorylation (OXPHOS) through the electron transport chain (ETC)^17,51^. The large GTPase, OPA1, further contributes to cristae architecture by stabilizing MICOS complex integrity^52^. Specifically, the long isoform of OPA1 (OPA1-L) mediates IMM fusion^53^ and maintains cristae lumen width, while the short isoforms (OPA1-S1/2) cooperatively regulate cristae lumen length^54^.

**Figure 3.**
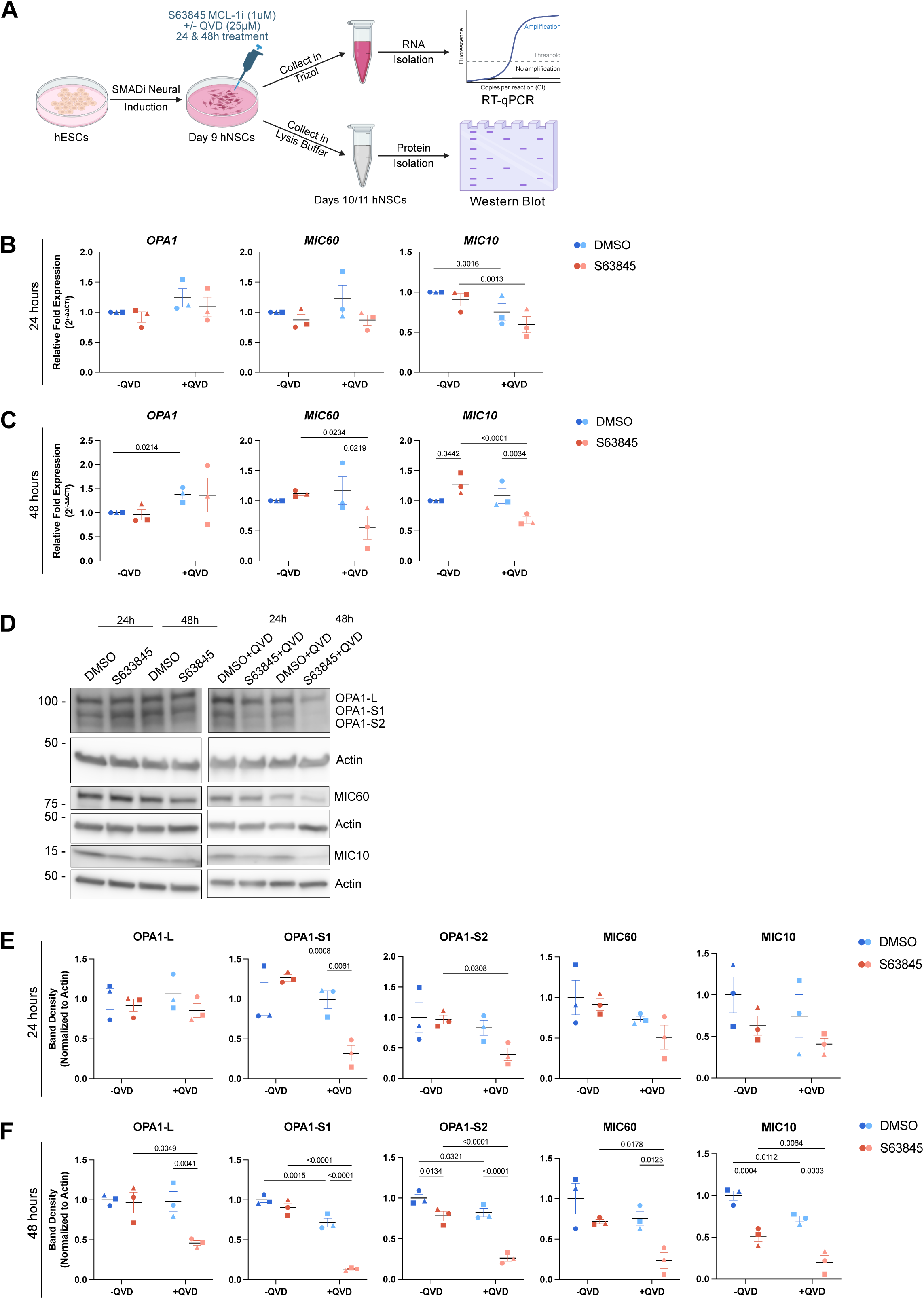
S63845 and QVD co-treatment reveal altered levels of cristae-shaping proteins. (**A**) Schematic detailing the neural induction of human embryonic stem cells (hESCs) using SMAD inhibition (SMADi) for the derivation of the human neural progenitor cells (NPCs) tissue culture model. NPCs were maintained in neural induction media (NIM) with daily media changes until they were passaged on day 7. NPCs were given a 24-hour acclimation period on day 8 before treatment on day 9 with 1µm S63845 (MCL-1 inhibitor) −/+ 25µM QVD (caspase inhibitor). Cells were then collected at 24- and 48-hours post treatment on days 10 and 11, respectively, in either Trizol for RNA extraction and real-time quantitative polymerase chain reaction (RT-qPCR) or protease inhibitor lysis buffer for protein isolation and western blot. (**B**) Quantified relative expression of *OPA1*, *MIC60*, and *MIC10* after 24h (top panel) and 48h (bottom panel) treatments. Two technical replicates were used for all RT- qPCR data. RT-qPCR data are displayed as relative fold expression derived from 2^(-ΔΔCT)^, and raw data were analyzed using ANCOVA as described in the methods section. (**C**) Representative western blot images (n=3) of OPA1-L, OPA1-S1, OPA1-S2, MIC60, and MIC10 after 24h and 48h treatments. (**D**) Quantified band density normalized to Actin loading control derived from western blots of OPA1-L, OPA1-S1, OPA1-S2, MIC60, and MIC10 after 24 (top panel) and 48h (bottom panel) treatments. Each data point represents a biological replicate. Shape of the data points corresponds to a biological replicate (circle for n=1, square for n=2, and triangle for n=3). Band density data were analyzed using an ordinary two-way ANOVA with an uncorrected Fisher’s LSD test for multiple comparisons. All tests used a 95% confidence level (p-value ≤ 0.05) as the significance threshold. Error bars represent standard error of the mean.

To determine whether MCL-1 inhibition affects the transcriptional regulation of these structural components, mRNA expression of *OPA1*, *MIC60*, and *MIC10* was assessed at 24 and 48 hours following S63845 treatment (**Figure 3B-C**). At 24 hours, MCL-1 inhibition did not significantly alter the expression of *OPA1*, *MIC60*, or *MIC10* (**Figure 3B**). At 48 hours, however, *MIC10* mRNA levels were significantly increased by MCL-1 inhibition, while *MIC60* and *OPA1* expression remained unchanged (**Figure 3C**). Co-treatment with the pan-caspase inhibitor QVD captured a reduction in the expression of both *MIC60* and *MIC10* at 48h, suggesting a caspase-dependent component to the transcriptional regulation of MICOS subunits under conditions of MCL-1 inhibition.

To determine whether the transcriptional changes observed at the mRNA level were reflected at the protein level, we next performed immunoblotting for OPA1 isoforms, MIC60, and MIC10 across all treatment conditions and timepoints (**Figure 3D**). At 24 hours, MCL-1 inhibition alone did not significantly alter OPA1 isoform levels; however, combined MCL-1 and caspase inhibition induced a significant reduction in OPA1-S1/2 protein levels (**Figure 3E**). At 48 hours, a reduction in OPA1-S1 and OPA1-S2 was observed across multiple conditions, with the most pronounced decrease occurring under combined S63845 and QVD co-treatment, while OPA1-L levels were also significantly reduced at this timepoint (**Figure 3F**). These isoform-specific reductions suggest that MCL-1 inhibition disrupts the stoichiometric balance between fusion-competent long-form OPA1 (OPA1-L) and its short isoforms (OPA1-S1/2), potentially reflecting an impairment of OPA1-L proteolytic processing rather than enhanced cleavage activity.

Assessment of MICOS core subunit abundance revealed no significant changes in MIC60 or MIC10 protein levels at 24 hours under any condition (**Figure 3E**). However, MIC13, which serves as a bridge between MIC60 and MIC10 subcomplex, was significantly reduced at 24 hours (**Supplementary Figure 3A-B**). At 48 hours, MCL-1 inhibition alone induced a significant reduction in MIC10 protein levels, despite transcriptional upregulation observed at this timepoint (**Figure 3C**), while MIC60 levels remained unchanged (**Figure 3F**). MIC25 and MIC13 were also destabilized following 48 hours S63845 treatment (**Supplementary Figure 3C**). At 48 hours, co-treatment with QVD further revealed a reduction in MIC60, MIC10, MIC19, MIC25, and MIC13 protein levels at 48 hours (**Figure 3F and Supplementary Figure 3C**), indicating that caspase inhibition unmasks a broader destabilization of the MICOS complex. Collectively, these findings identify MCL-1 as a critical upstream regulator of mitochondrial inner membrane organization, acting through parallel destabilization of OPA1 isoform homeostasis and MICOS complex integrity to maintain cristae architecture.

To assess the consequences of these mitochondrial changes on cellular respiration, we performed respirometry assays using an Agilent Seahorse XF Mito Stress Test to measure oxygen consumption rate (OCR). At 24h of MCL-1 inhibition, there was a significant decrease in maximal respiration, spare respiratory capacity, and the percentage of spare respiratory capacity, which were not rescued by QVD (**Supplementary Figure 4A-B**). The failure of QVD to rescue any of the observed OCR deficits confirms that the respiratory impairment is not secondary to caspase activation or apoptotic mitochondrial permeabilization. Co-treatment with QVD further revealed decreased coupling efficiency and non-mitochondrial OCR (**Supplementary Figure 4A-B**). At 48h of MCL-1 inhibition, the percentage of spare respiratory capacity remained decreased and was not rescued by QVD. Co-treatment with QVD at 48h revealed a decrease in basal respiration, maximal respiration, ATP-linked production, non-mitochondrial OCR, and spare respiratory capacity (**Supplementary Figure 4C-D**). Thus, cristae changes induced by MCL-1 inhibition cause a progressive, caspase- independent impairment of mitochondrial respiration, characterized by deficits in maximal respiration and spare respiratory capacity at 24 hours that worsen by 48 hours to include reductions in basal respiration and ATP-linked production.

### MCL-1 inhibition impairs fatty acid oxidation (FAO)

We predicted that FAO transmembrane enzymes and transport proteins rely heavily on the proper assembly and structure of the OMM and IMM for proper activity (**Figure 4A**). Previous investigations have shown close interactions between MCL-1 and FAO enzymes in cancer cells^55^, and mouse studies have also established the importance of FAO for the maintenance of embryonic neurodevelopment and adult neural stem cells^56^. To explore whether the mitochondrial structural perturbations observed upon MCL-1 inhibition extend to mitochondrial metabolic function, we assessed key regulators of fatty acid β-oxidation (**Figure 4B**). At 24 hours, CPT1a protein levels were increased following MCL-1 inhibition, while ACSL1, CPT2, and VLCAD levels were not significantly affected (**Figure 4C**). Co-treatment with QVD at 24 hours revealed a significant decrease of CPT1a, CPT2, and VLCAD levels compared to DMSO, with ACSL1 also showing a reduction, albeit not significant compared to vehicle control, but significantly reduced when compared to MCL-1 alone (**Figure 4C**). At 48 hours of MCL-1 inhibition alone, CPT1a protein levels remained elevated along with VLCAD, while CPT2 levels significantly decreased (**Figure 4D**). Co-treatment with QVD at 48 hours resulted in higher levels of CPT1a and a significant decrease in CPT2 and VLCAD compared to control (**Figure 4D**).

**Figure 4.**
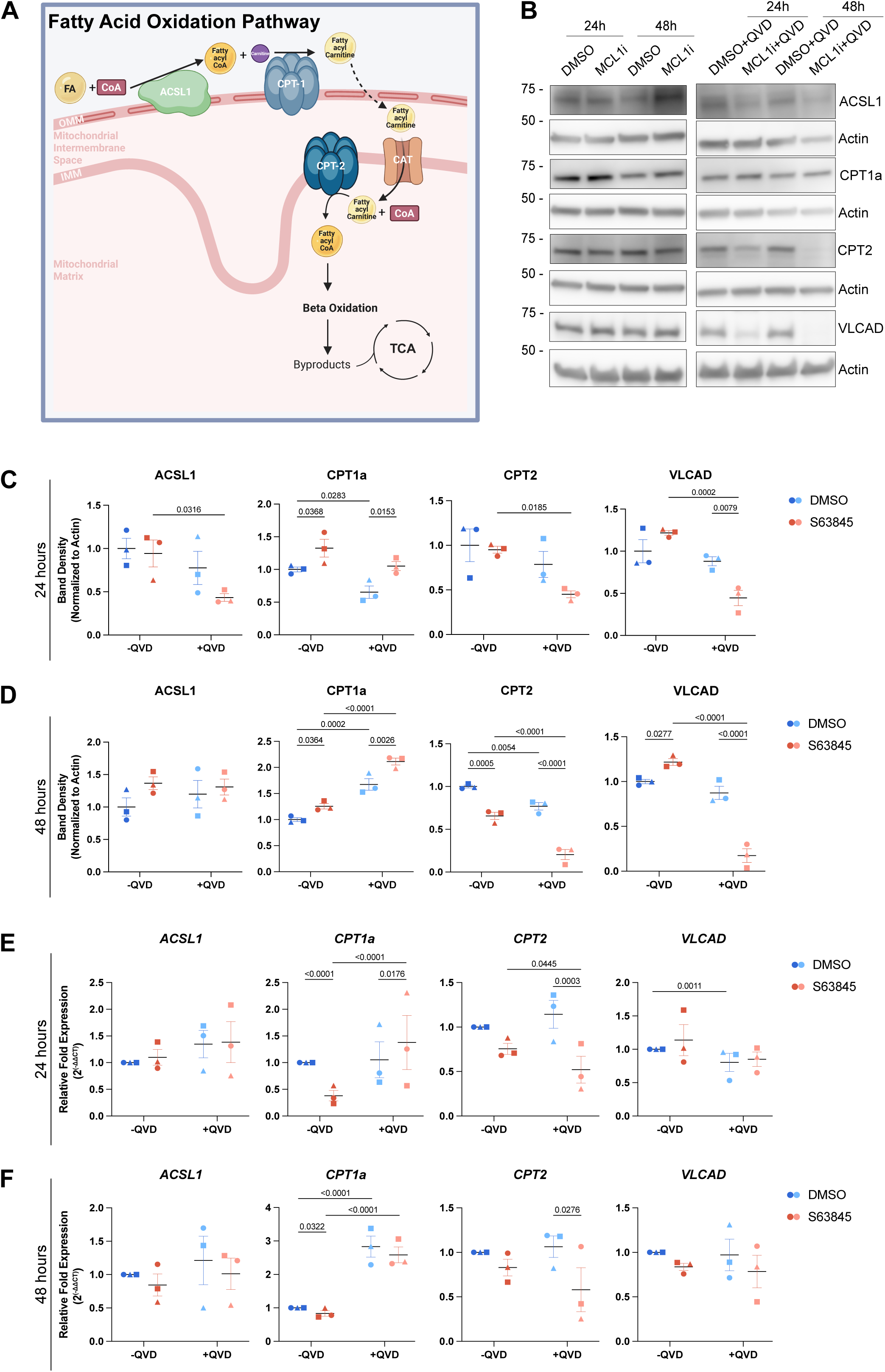
S63845 and QVD co-treatment results in downregulation of carnitine palmitoyltransferase and altered levels of FAO enzymes. (**A**) Schematic of the mitochondrial fatty acid beta-oxidation pathway. (**B**) Representative western blot images (n=3) and (**C**) quantified band density normalized to Actin loading control of FAO enzymes ACSL1, CPT1a, CPT2, and VLCAD after S63845 (MCL-1 inhibitor) −/+ QVD (caspase inhibitor) treatments for 24 hours (top panel) and 48 hours (bottom panel). Band density data were analyzed using an ordinary two-way ANOVA with an uncorrected Fisher’s LSD test for multiple comparisons. (**D**) Quantified relative expression of FAO enzymes *ACSL1*, *CPT1a*, *CPT2*, and *VLCAD* after 24h (top panel) and 48h (bottom panel) treatments. Two technical replicates were used for all RT-qPCR data. RT-qPCR data are displayed as relative fold expression derived from 2^(-ΔΔCT)^, and raw data were analyzed using ANCOVA as described in the methods section. Each data point on the graph represents a biological replicate. Shape of data point corresponds to each biological replicate (circle for n=1, square for n=2, and triangle for n=3). All tests used a 95% confidence level (p-value ≤ 0.05) as the significance threshold. Error bars represent standard error of the mean.

Previous studies reported a global downregulation of the FAO pathway at the mRNA level upon MCL-1 deletion in B-cell acute lymphoblastic leukemia (B-ALL cells)^25^. We performed RT-qPCR-based mRNA expression analyses to examine the expression patterns of FAO genes in NPCs treated with S63845. At 24 and 48 hours of MCL-1 inhibition alone, *ACSL1*, *CPT2*, and *VLCAD* levels remained unchanged while *CPT1a* was significantly downregulated (**Figures 4E-F**) despite higher protein levels detected at both timepoints (**Figure 4C-D**). This suggests that CPT1a protein accumulation occurs through a post-transcriptional mechanism that stabilizes the protein under combined MCL-1 and caspase inhibition rather than transcriptional upregulation. This relationship between in *CPT1a* mRNA and protein persists in the presence of QVD at 48h (**Figure 4F**). At 24 hours, however, inhibition of MCL-1 produced a significant upregulation of *CPT1a* mRNA, albeit with high variability (**Figure 4E**). QVD co-treatment resulted in upregulation of *CPT2* at both timepoints (**Figures 4E-F**). Notably, this transcriptional decrease in *CPT2* was accompanied by a corresponding reduction in CPT2 protein (**Figure 4B-D**), suggesting that caspase inhibition engages an additional transcriptional suppression of CPT2 beyond the post-transcriptional effects seen with MCL-1 inhibition alone. Consistent with the transcriptional suppression previously reported^25^, we observe downregulation of *CPT1a* mRNA and protein with MCL-1 inhibition alone, while other genes encoding for FAO enzymes, including ACSL1, CPT2, and VLCAD, show no transcriptional change despite protein-level alterations. This discordance between mRNA and protein levels in our model points to post-translational mechanisms as the primary drivers of FAO enzyme loss, rather than the transcriptional downregulation previously reported upon Mcl-1 deletion in cancer cells.

We next sought to determine whether enzyme dysregulation impairs FAO by directly assessing cellular fatty acid storage. Fatty acids get stored in lipid droplet organelles when they are not processed by β-oxidation. Accumulation of lipid droplets upon MCL-1 inhibition was previously reported in cancer cells and murine liver as a marker of disrupted fatty acid β- oxidation^26^. To further investigate the impact of MCL-1 inhibition on FAO, NPCs were cultured in nutrient-deficient DMEM media supplemented only with palmitate during treatment with MCL-1 inhibitor for 12 hours, followed by a 2-hour starvation period in HBSS under continuous MCL-1 inhibition with S63845 (**Figure 5A**). Consistent with altered FAO enzyme levels detected by RT-qPCR and immunoblotting and previous reports, S63845-treated NPCs accumulated excess neutral lipids when challenged with palmitate as their only resource for fuel (**Figure 5B**). Quantitative image analysis revealed a significant increase in average lipid droplet volume and a trending increase in maximum volume (**Figure 5C**), indicative of reduced lipid turnover and a shift toward storage. These data suggest that MCL-1 sustains metabolic flux as fatty acids fail to import to the matrix and accumulate in lipid droplets.

**Figure 5.**
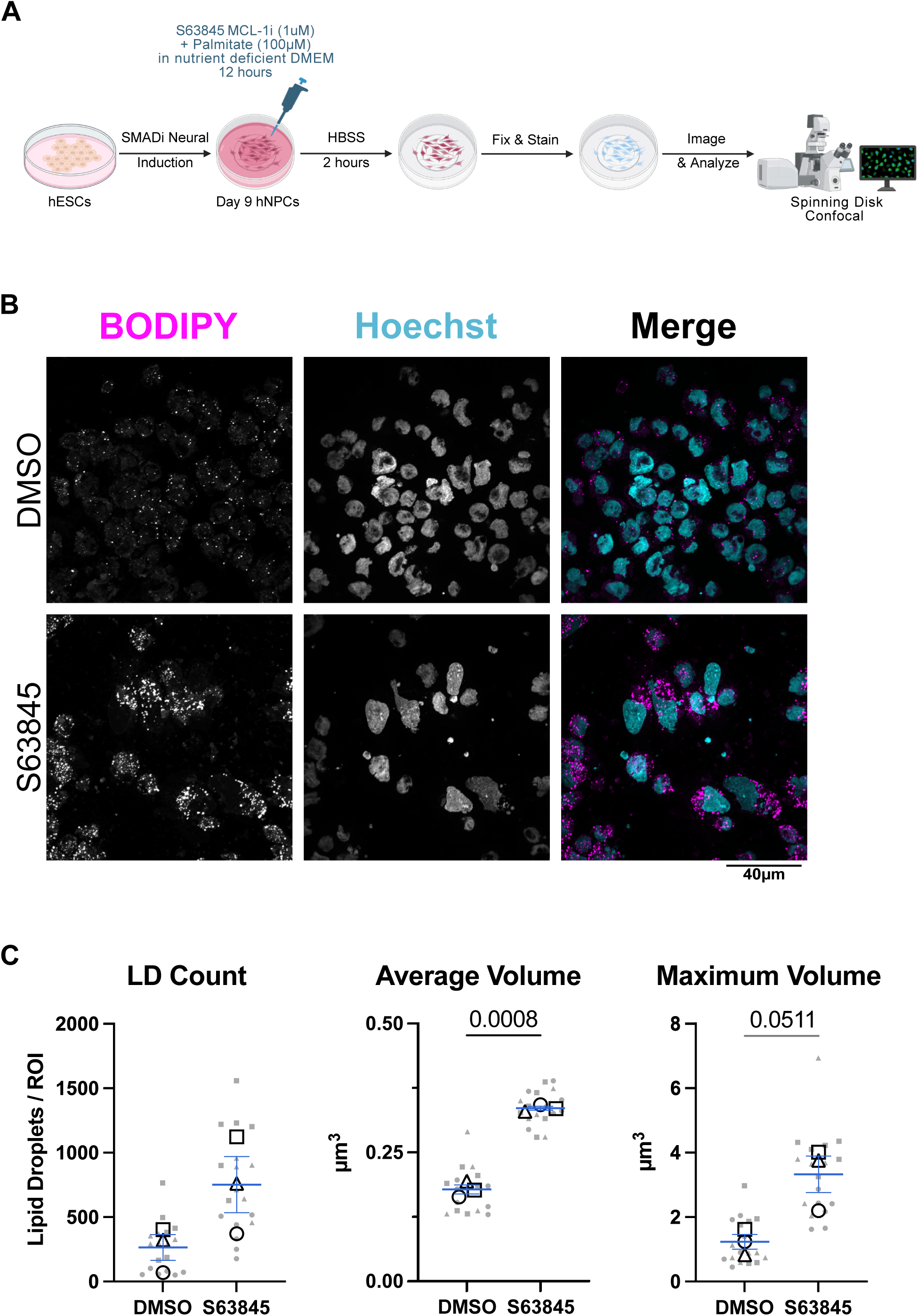
NPCs supplemented with palmitate and treated with S63845 accumulate lipid droplets. (**A**) Schematic detailing the neural induction of human embryonic stem cells (hESCs) using SMAD inhibition (SMADi) for the derivation of the human neural progenitor cells (NPCs) tissue culture model. NPCs were maintained in neural induction media (NIM) with daily media changes until they are passaged on day 7. NPCs were given a 24-hour acclimation period on day 8 before treatment on day 9 with 1µm S63845 (MCL-1 inhibitor) in nutrient deficient DMEM media (-glucose, -glutamine, -pyruvate) and supplemented with 100µM of palmitate for 12 hours followed by a 2-hour starvation in HBSS. Cells were then live-stained with BODIPY and Hoechst prior to paraformaldehyde fixation for imaging on an SDC (Spinning Disk Confocal) microscope. (**B**) Representative immunofluorescent images (n=3) of lipid droplets (BODIPY in magenta) and nuclei (Hoechst in cyan) were acquired on a spinning disk confocal microscope at 100X (scale bar = 40µm). (**C**) Quantification of lipid droplets per image, average volume, and maximum volume. Each large, opened-shape data point (black) represents the mean of a biological replicate, and each small, closed-shape data point (transparent gray) represents a technical replicate (a single image). Shape of the data points corresponds to a biological replicate (circle for n=1, square for n=2, and triangle for n=3). Data were analyzed using an ordinary two-way ANOVA with an uncorrected Fisher’s LSD test for multiple comparisons and a 95% confidence level (p-value ≤ 0.05 as the significance threshold). Error bars represent standard error of the mean.

### MCL-1 inhibition impairs FAO by delocalizing ACLS1 from the outer mitochondrial membrane

Previous studies in cancer cells identified an ACSL1 BH3-like binding domain that mediates a direct interaction between ACSL1 and MCL-1 at the outer mitochondrial membrane (OMM)^23,26^. Given that ACSL1 catalyzes the first committed step of the FAO cascade, its proper localization to the OMM is essential for efficient fatty acid activation and subsequent mitochondrial import. To investigate whether MCL-1 serves as a scaffold for ACSL1 recruitment or retention at the OMM, we generated a stable hESC line overexpressing ACSL1:HaloTag and differentiated these cells into NPCs for live-cell imaging. ACSL1:HaloTag was detected using a JFX dye and mitochondria were labeled with MitoTracker Green FM. To achieve super-resolution, cells were live-imaged on a SoRa SDC using a 100X objective with 2.8X magnification (**Figure 6A**). Co-localization image analysis revealed a significant decrease in mitochondria-associated ACSL1 under two defining parameters following MCL-1 inhibition at both 24 and 48 hours (**Figure 6B**), indicating that MCL-1 is required for the proper retention of ACSL1 at the OMM. Together with the lipid droplet accumulation data, these findings suggest that MCL-1 inhibition disrupts FAO at its earliest enzymatic step by displacing ACSL1 from its site of action, thereby reducing fatty acid activation and mitochondrial import capacity.

**Figure 6.**
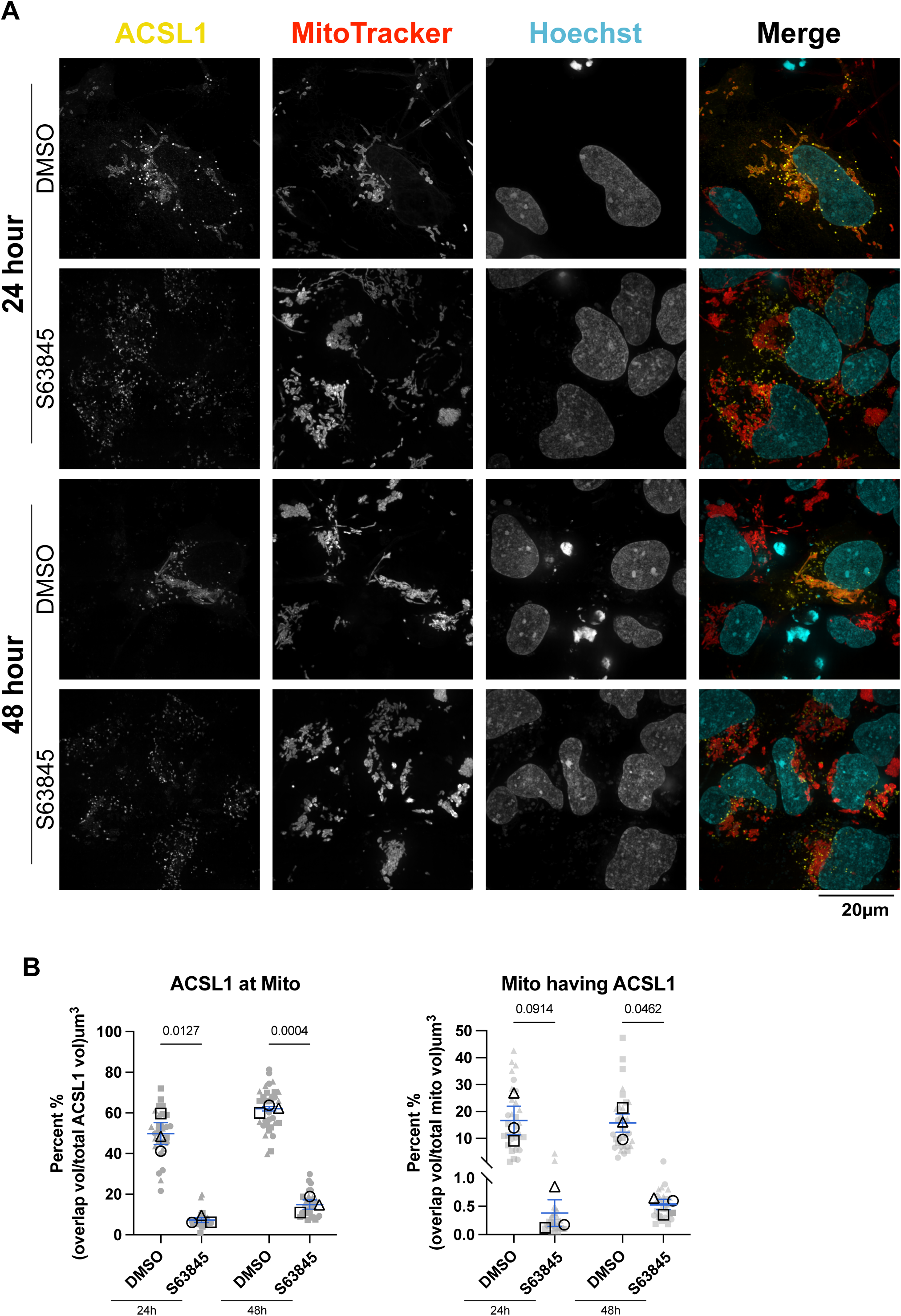
S63845 displaces ACSL1 from the mitochondria. (**A**) Immunofluorescent images of ACSL1:HaloTag (JFX in yellow) mitochondria (MitoTracker Green FM in red) and nuclei (Hoechst in cyan) were acquired live on SoRa SDC (Super Resolution by Optical Reassignment Spinning Disk Confocal) microscope at 100X with 2.8X magnification (scale bar = 20µm). Representative images (n=3) of NPCs treated with S63845 (MCL-1 inhibitor) for 24- (top panel) and 48-hours (bottom panel). (**B**) Quantification of total ACSL1 localized at the mitochondria (left panel) and calculated percentage of mitochondria with ACSL1 instances (right panel). Each large, opened-shape data point (black) represents the mean of a biological replicate, and each small, closed-shape data point (transparent gray) represents a technical replicate (a single image). Shape of the data points corresponds to a biological replicate (circle for n=1, square for n=2, and triangle for n=3). Data were analyzed using an ordinary two-way ANOVA with an uncorrected Fisher’s LSD test for multiple comparisons and a 95% confidence level (p-value ≤ 0.05 as the significance threshold). Error bars represent standard error of the mean.

### MCL-1 maintains neural progenitor identity and supports neuronal differentiation

Given the role of MCL-1 in FAO and the mitochondrial and metabolic perturbations triggered by MCL-1 inhibition, we investigated the effects of S63845 treatment on neurogenic progression. All findings were observed on differentiation days 10 and 11 for 24- and 48-hour treatments, respectively. Although we did not detect changes in global protein levels of NPC identity markers (**Supplementary Figure 5A-C**), we observed a significant downregulation of various identity markers (**Supplementary Figure 5D-E**). At 24h of MCL-1 inhibition alone, *PAX6* (a marker of neural stem cell identity), *EOMES* (the gene encoding TBR2, a marker of intermediate progenitor cells (IPCs)), *TBR1* (a postmitotic marker of newborn glutamatergic projection neurons), and *TUBB3* (neuronal microtubule and early marker for newborn neurons) were downregulated (**Supplementary Figure 5D**). Co-treatment with QVD at 24h rescued *PAX6* and *EOMES* mRNA levels, but not *TBR1* and *TUBB3* (**Supplementary Figure 5D**). At 48h of MCL-1 inhibition alone, *PAX6*, *EOMES*, and *TBR1* remained downregulated, but not *TUBB3* (**Supplementary Figure 5E**). Co-treatment with QVD at 48h persistently rescued *PAX6* as well as *TBR1* but not *EOMES* and maintained a downregulation in *TUBB3* (**Supplementary Figure 5E**). We followed these analyses with immunofluorescent imaging, which revealed a small population of IPCs (PAX6^-^/TBR2^+^) in the culture of NPCs. IPCs are progenitor cells that are committed to give rise to deep layer glutamatergic projection neurons. MCL-1 inhibition selectively reduced the proportion of IPCs at 24 hours of MCL-1 inhibition without rescue by QVD co-treatment (**Figure 7A**). Loss of IPCs persisted at 48 hours (**Figure 7B**). These findings suggest that MCL-1 is critical for maintaining the identity of this progenitor pool through non-apoptotic mechanisms. To determine how these effects on IPCs affect neuronal differentiation, we first examined cell proliferation using EdU labeling and found no significant changes in early or late S-phase or in global proliferation states across all treatment conditions and timepoints (**Supplementary Figure 6A-D**). To further investigate the potency of neural stem/progenitor cells, we examined their ability to differentiate and give rise to newborn neurons using βIII-tubulin staining (**Figures 8A-B**). Control cells formed dense neurite networks with extensive βIII-tubulin labeling, whereas MCL-1-inhibited neurons displayed sparse and truncated processes, reflecting defective neurite extension and maturation at both timepoints (**Figure 8A-B**). When treated with the MCL-1 inhibitor, neural stem/progenitor cells give rise to fewer newborn neurons with retracted neuronal processes (**Figure 8C-D**); however, most parameters were rescued upon co-treatment with QVD, except for average major axis length and average surface area at 48h of S63845 treatment. Taken together, these findings shed light on the importance of MCL-1 for the potency of neural stem/progenitor cells to undergo proper differentiation and for newborn neurons to undergo proper maturation. The absence of overt cytotoxicity suggests that MCL-1 directly supports neuronal commitment rather than survival alone.

**Figure 7.**
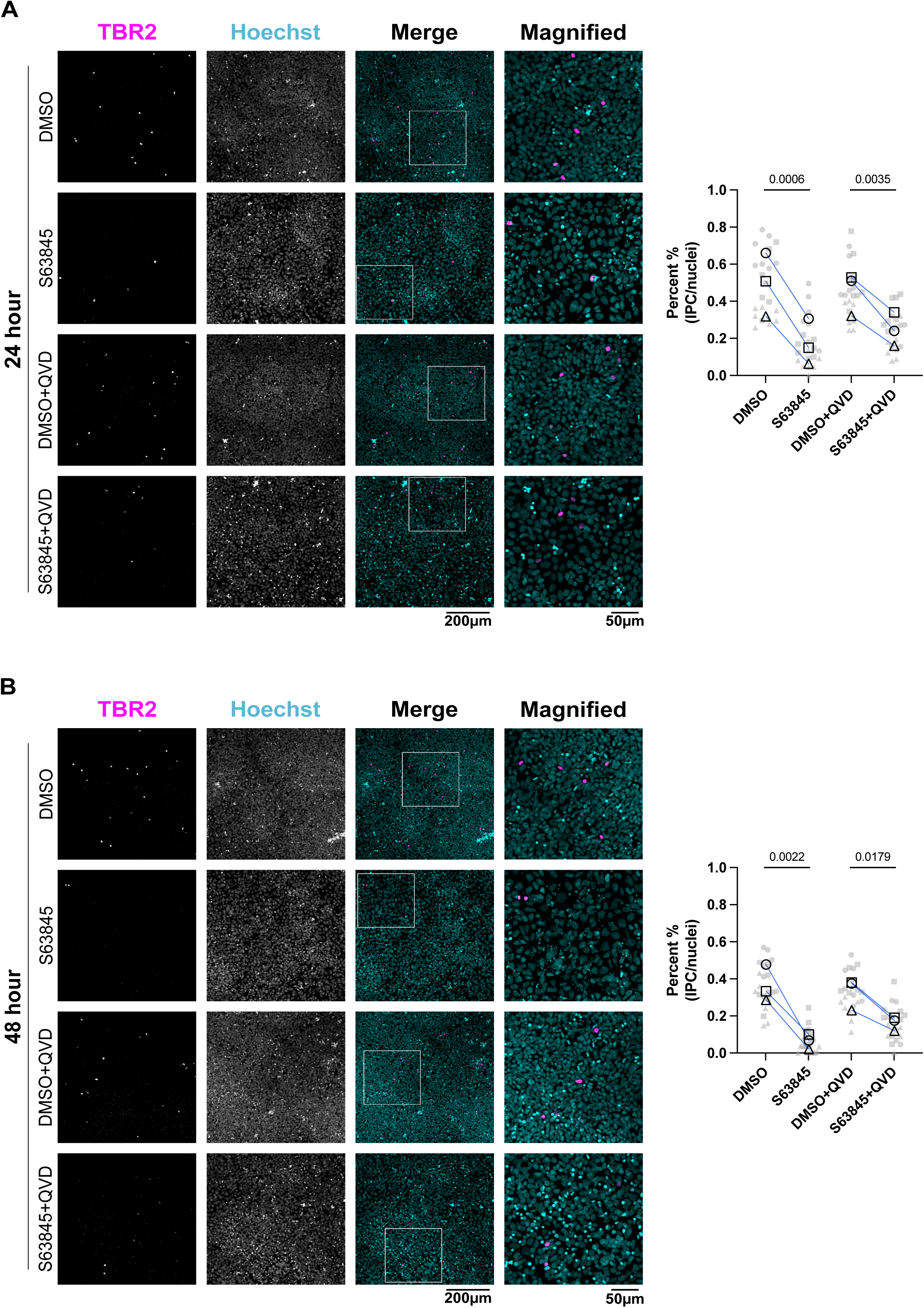
S63845 results in depletion of the TBR2^+^/PAX6^-^ IPC pool. Immunofluorescent images of TBR2 (in magenta) and nuclei (Hoechst in cyan) were acquired on SDC (Spinning Disk Confocal) at 20X (scale bar = 200µm) and magnified panels (scale bar = 50µm). Representative images (n=3) of NPCs treated with S63845 (MCL-1 inhibitor) −/+ QVD (caspase inhibitor) for 24- (**A**) and 48-hours (**B**). (**A-B**) Blind count of TBR2^+^ IPCs calculated as percentage of total cell population. Each large, opened-shape data point (black) represents the mean of a biological replicate, and each small, closed-shape data point (transparent gray) represents a technical replicate (a single image). Shape of the data points corresponds to a biological replicate (circle for n=1, square for n=2, and triangle for n=3). Data were analyzed using a repeated measures two-way ANOVA with an uncorrected Fisher’s LSD test for multiple comparisons and a 95% confidence level (p-value ≤ 0.05 as the significance threshold). Error bars represent standard error of the mean.

**Figure 8.**
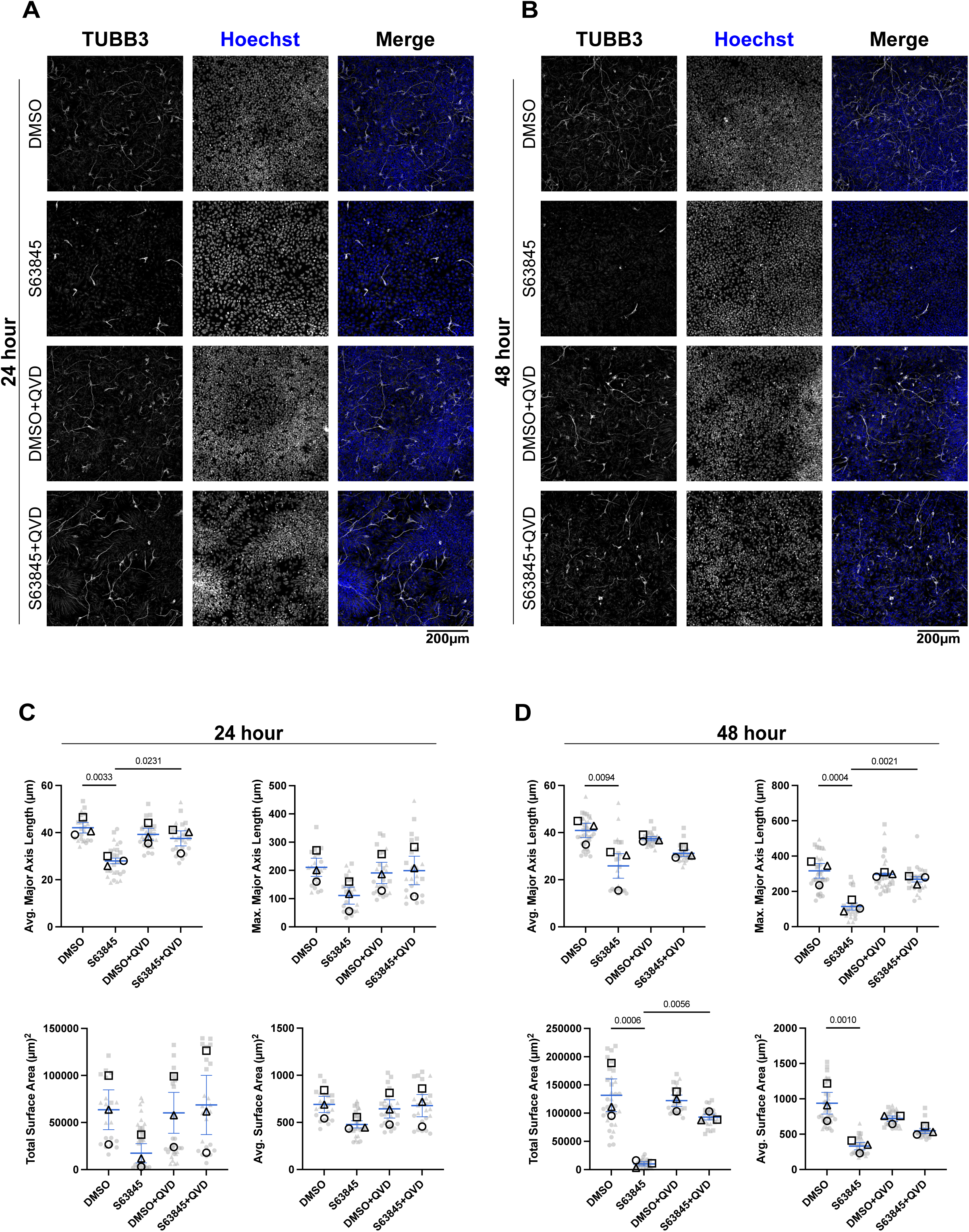
S63845 results in the depletion of newborn neurons detected by BIII-Tubulin. Immunofluorescent images of TUBB3 (in white) and nuclei (Hoechst in blue) were acquired on SDC (Spinning Disk Confocal) at 20X (scale bar = 200µm). Representative images (n=3) of NPCs treated with S63845 (MCL-1 inhibitor) −/+ QVD (caspase inhibitor) for 24- (**A**) and 48- hours (**B**). Quantified morphology analysis of TUBB3 parameters, including average and maximum major axis length and total and average surface area after 24h (**C**) and 48h (**D**) treatments. Each large, opened-shape data point (black) represents the mean of a biological replicate, and each small, closed-shape data point (transparent gray) represents a technical replicate (a single image). Shape of the data points corresponds to a biological replicate (circle for n=1, square for n=2, and triangle for n=3). Data were analyzed using an ordinary two-way ANOVA with an uncorrected Fisher’s LSD test for multiple comparisons and a 95% confidence level (p-value ≤ 0.05 as the significance threshold). Error bars represent the standard error of the mean.

## Discussion

The non-apoptotic functions of MCL-1 in maintaining mitochondrial homeostasis are increasingly recognized across diverse cell types^18,26,27,32^, yet their relevance to human neurodevelopment has remained largely unexplored. Our findings establish MCL-1 as a critical coordinator of mitochondrial architecture and metabolic competence in human neural progenitor cells, extending this paradigm to the developing human brain. By integrating ultrastructural, molecular, and functional analyses, we demonstrate that MCL-1 sustains cristae integrity through stabilization of OPA1 isoform homeostasis and the MICOS complex, while simultaneously retaining ACSL1 at the OMM to enable fatty acid activation. Disruption of either arm of this coordinated function impairs FAO metabolism, reduces mitochondrial oxygen consumption, and ultimately compromises the identity of intermediate progenitors and neuronal maturation. Together, these findings reposition MCL-1 from a canonical anti-apoptotic protein to a structural and metabolic molecule that is indispensable for proper human neurogenesis.

While MCL-1 has been reported to influence FAO in cancer cells and to localize to the IMM, its impact on cristae integrity in neural progenitors remained unexplored. Our findings bridge this gap by demonstrating that MCL-1 is necessary for maintaining cristae organization and FAO competence in NPCs. The observed mitochondrial morphological swelling upon MCL-1 inhibition likely reflect a compensatory response to the loss of metabolic function, as evidenced by lipid droplet accumulation and impaired oxygen consumption. We find this disruption to be coupled to a decline in FAO enzymes. Consistent with findings in cancer cells, we demonstrate that MCL-1 inhibition causes displacement of ACSL1 from the OMM in NPCs. Given that all altered FAO enzyme levels were downstream of ACSL1, this raises the question of whether ACSL1 represents a rate-limiting step in the mitochondrial fatty acid oxidation pathway in neural progenitor cells. If ACSL1 is displaced from the mitochondria, this may disrupt a feedback loop that maintains the expression and stabilization of downstream FAO enzymes, thereby dismantling the fatty acid oxidation cascade and driving lipid accumulation. The convergence of ACSL1 delocalization with cristae structural defects suggests that MCL- 1 coordinates fatty acid activation at the OMM and oxidative capacity at the IMM through spatially distinct but functionally coupled mechanisms.

Beyond the broad transcriptional suppression of FAO genes previously reported upon MCL-1 deletion in cancer cells^25^, our data indicate post-translational mechanisms, including altered protein stability, as the primary drivers of FAO enzyme dysregulation in NPCs. This distinction is conceptually important: it suggests that in neural progenitors, MCL-1 supports FAO enzyme function primarily through structural maintenance of the mitochondrial membrane environment rather than through transcriptional control and raises the possibility that the cristae collapse further impairs the stability and localization of membrane-associated FAO enzymes.

We propose a model in which MCL-1 operates through two coordinated and caspase-independent mechanisms to sustain mitochondrial function in NPCs. At the OMM, MCL-1 retains ACSL1 to enable fatty acid activation upstream of CPT1/2, consistent with the direct BH3-like interaction between MCL-1 and ACSL1 previously identified in cancer cells. At the IMM, MCL-1 preserves cristae integrity through stabilization of OPA1 isoform stoichiometry and the MICOS complex, maintaining the structural platform required for OXPHOS and FAO enzyme function. The combined loss of OMM-level fatty acid activation and IMM-level cristae integrity and respiration infer that these functions operate in parallel and require the non-apoptotic capacities of MCL-1. Distinguishing the direct OMM scaffolding role of MCL-1 for ACSL1 from potential indirect effects of cristae collapse on ACSL1 retention will require future studies using structure-specific MCL-1 mutants or domain-selective approaches.

The metabolic intermediates generated by FAO, particularly acetyl-CoA, are well-established regulators of chromatin state through their role as substrates for histone acetyltransferases. A reduction in mitochondrial acetyl-CoA availability secondary to FAO impairment could alter histone acetylation at neurogenic loci, thereby influencing transcriptional programs that govern progenitor identity and lineage commitment. This metabolic-epigenetic axis represents a plausible mechanism by which MCL-1-dependent FAO supports the maintenance of intermediate progenitors and constitutes a testable model for future investigation. In the context of neurodevelopment, where bioenergetically demanding progenitor cells must maintain a precise metabolic profile, mitochondrial competence functions as a quality-control checkpoint for lineage progression.

The clinical precedent established by venetoclax, a selective BCL-2 inhibitor approved for hematologic malignancies^57,58^, underscores both the therapeutic potential and the selectivity challenges inherent to targeting BCL-2 family members. The potential cardiotoxicity and emerging neural concerns associated with MCL-1 inhibitors, relative to the comparatively favorable CNS tolerability profile of venetoclax, reinforce the notion that BH3-mimetic selectivity can be influenced by tissue-specific factors. Our data suggest that cells undergoing active neurogenesis are particularly vulnerable to MCL-1 inhibition because they rely on MCL- 1’s non-apoptotic roles rather than its pro-survival function, a distinction that the clinical experience with venetoclax does not capture and that could be critical when evaluating MCL- 1-targeting agents in oncology regimens.

The contribution of BCL-XL to mitochondrial bioenergetics provides an important parallel for interpreting our findings. BCL-XL has been shown to interact directly with subunits of the electron transport chain (ETC), particularly complexes I and III, enhancing proton flux efficiency and increasing ATP synthesis independently of its anti-apoptotic function^59,60^. These studies established the precedent that BCL-2 family members can act as direct modulators of ETC stoichiometry and complex assembly, factors that MCL-1 activity in the IMM could also modulate. Revealing the contribution of MCL-1 to these critical aspects of mitochondrial function will require profiling respiratory complex assembly under MCL-1 inhibition to help clarify whether impaired respiration in NPCs is also due to direct effects on the ETC.

The mechanism by which MCL-1 inhibition destabilizes OPA1 isoform stoichiometry likely converges on the inner membrane proteases OMA1 and YME1L, which execute opposing cleavage events to regulate the balance between long and short OPA1 isoforms^61^. Under basal conditions, YME1L-mediated constitutive processing of OPA1 is balanced against stress-induced OMA1 activation, which rapidly converts long OPA1 isoforms to short forms and thereby promotes cristae widening and cytochrome c release competence. Mitochondrial membrane depolarization and matrix stress are well-established triggers for OMA1 hyperactivation. The metabolic impairment and FAO failure we observe upon MCL-1 inhibition could create an environment conducive to OMA1 activation, shifting the OPA1 equilibrium toward short isoforms and destabilizing the cristae junctions that MCL-1 normally maintains. MCL-1 is likely targeted for degradation by YME1L^49^ (Perciavalle, 2012), but whether OPA1 cleavage imbalance is a downstream consequence of the broader metabolic failure we describe remains an important open question for future mechanistic studies.

The influence of mitochondrial architecture on developmental fate decisions is increasingly recognized. *In vivo* and *in vitro* studies have identified a postmitotic window of neurogenic plasticity controlled by mitochondrial morphology^16^, and that disruption of the FAO pathway promotes asymmetric differentiation of adult mouse pluripotent and neural stem cells^37,43^, underscoring the importance of mitochondrial morphology and FAO in stem cell fate regulation. Our data reveal that MCL-1 inhibition disrupts this balance through mechanisms extending beyond FAO itself, implicating cristae architecture and ACSL1-mediated lipid activation as additional determinants of progenitor identity. Importantly, mutations in cristae-shaping proteins, including OPA1 and MICOS subunits, are associated with severe neurodevelopmental and neurodegenerative disorders such as dominant optic atrophy, and Parkinson’s disease-associated neurodegeneration. The mechanistic parallels between these genetic disorders and the phenotypes induced by MCL-1 inhibition in NPCs reinforce the notion that mitochondrial function is indispensable for these bioenergetically demanding neural cells.

Taken together, our results position MCL-1 upstream of OPA1, MICOS, and ACSL1, and define a previously unappreciated dimension of MCL-1 function in human neurogenesis. MCL-1 is emerging as a metabolic modulator coordinating cristae architecture, ACSL1 mitochondrial localization, lipid oxidation, mitochondrial respiration, and lineage progression, all independently of its canonical anti-apoptotic role. These findings carry potential translational relevance. MCL-1 inhibitors have been explored as therapeutic agents for chemoresistant malignancies, but clinical development has been limited by cardiotoxicity. We previously reported that MCL-1 compromises cytoskeletal integrity in cardiomyocytes as detected by phalloidin, and here we report failed maturation of newborn neurons as detected by cytoskeletal marker, TUBB3. Our data suggest that the non-apoptotic consequences of MCL-1 inhibition in neural progenitors require consideration when evaluating the neurodevelopmental safety of BH3 mimetic therapies, particularly in pediatric contexts when active neurogenesis is ongoing. Further investigation into how MCL-1 coordinates cristae organization and lipid metabolism in progenitor cells may illuminate new vulnerabilities in neurodevelopmental and neurodegenerative disorders.

### Limitations

The use of caspase inhibitors to assess the involvement of apoptosis is a limitation of our study; ideally, apoptosis would be more specifically blocked by targeting BAX/BAK. However, these cells exhibit dramatic changes in mitochondrial morphology that would confound results^62^. We rationalized that phenotypes detected with S63845 alone, but not S63845 co-treatment with QVD, are rescued by inhibition of apoptosis, suggesting they are attributable to MCL-1’s canonical anti-apoptotic functions. Conversely, phenotypes detected with S63845 and QVD co-treatment, but not with S63845 alone, likely represent a snapshot of cellular changes occurring in vulnerable cells before they undergo apoptosis and are therefore also attributable to MCL-1’s canonical anti-apoptotic role. Finally, phenotypes detected under S63845 that persist with QVD co-treatment, thus not rescued by caspase inhibition, are attributed to MCL-1’s non-apoptotic functions.

In this study, we aimed to disrupt the BH3 binding domain of MCL-1 through pharmacological inhibition using S63845. The small molecular inhibitor, S63845, selectively occupies the BH3- binding groove of MCL-1, blocking its protein-protein interactions while preserving MCL-1 protein levels. This approach allowed us to specifically investigate the consequences of BH3- domain blockade in isolation from broader transcriptional and translational adaptations that accompany genetic depletion. As demonstrated by our immunoblot data, S63845 treatment increases total MCL-1 protein levels, consistent with stabilization of the protein when its BH3 domain is occupied. This approach also holds direct clinical relevance, as S63845 and related BH3 mimetics are pharmacological agents under therapeutic investigation, making the phenotypes described here translatable to the consequences of therapeutic MCL-1 inhibition on human neural cells. Nevertheless, we acknowledge that stabilized, but inhibited MCL-1 may retain partial scaffolding or structural functions at the IMM or OMM that would be absent in a true loss-of-function model. Future studies employing swap genetic tools^27^ or inducible genetic deletion of MCL-1 in NPCs will be valuable for distinguishing the consequences of BH3-domain blockade from the complete loss of MCL-1 protein, and for determining whether the mitochondrial structural and metabolic phenotypes described here are fully recapitulated under conditions of genetic loss of function. A further challenge in fully dissecting the distinct contributions of MCL-1 at the OMM and IMM is the lack of tools that selectively restrict MCL- 1 localization to one membrane compartment without perturbing the protein itself. Current approaches to enforce IMM-restricted localization rely on the addition of exogenous targeting domains, which may alter MCL-1 conformation, interaction partners, or function. The development of refined tools that enable compartment-specific interrogation of MCL-1 function without modifying the endogenous protein is an important priority for the field.

Lastly, our findings on the IPC pool depletion upon S63845 treatment strongly suggest a unique reliance on MCL-1. However, the differentiation protocols used to generate neural progenitor cultures *in vitro* yield a minority pool of TBR2-expressing IPCs that constitutes fewer than 1% of the total cell population. Prior to investigating the effects of S63845 on this cell type, we optimized the derivation of IPCs from various hPSC lines. Each study should identify the stage at which the TBR2 population reaches maximal expansion during hPSC differentiation to NPCs, as hESC and hiPSC lines exhibit variable competence in generating TBR2-positive intermediate progenitor cells, and the timing of peak generation varies across lines. These limitations highlight the need to develop methods to isolate and expand the intermediate progenitor cell population from hPSCs, which would be a valuable tool for future studies.

## Methods

### Human embryonic stem cells (hESCs)

Human embryonic stem cell line, H9 (WA09), was obtained from WiCell Research Institute (Wisconsin). H9s were maintained in mTeSR (STEMCELL Technologies, cat # 85850) media on 6-well plates coated with Matrigel (Corning, cat # 354277) at 37°C with 5% CO2. Culture medium was changed daily. Cells were checked daily for differentiation and were passaged every 3-4 days using Gentle cell dissociation solution (STEMCELL Technologies, cat # 07174). All experiments were performed under the supervision of the Vanderbilt Institutional Human Pluripotent Cell Research Oversight (VIHPCRO) Committee. Cells were periodically checked for contamination.

### hESC-derived NPC culture

H9s were cultured until reaching 70-80% confluency, ensuring minimal differentiation at the colony edges. Cells were dissociated using 1 mL of a Gentle Cell Dissociation Reagent (STEMCELL Technologies, cat # 100-0485) and incubated at 37°C for 8 minutes. 2 mL of DMEM-F12 (Thermo Scientific, cat # 11320033) supplemented with 10 µM of ROCK inhibitor (STEMCELL Technologies, cat # 72307) was added to each well. Cells were gently scraped, pipetted for transfer into a conical tube, and centrifuged at 300 x g for 5 minutes. Cell pellet was resuspended in STEMdiff™ SMADi <u>N</u>eural Induction <u>M</u>edium (NIM) (STEMCELL Technologies, cat # 08581) supplemented with 10 uM ROCK inhibitor. Cell suspension was counted and plated at a density of 2.0-2.5x10^6^ cells per well in a 6-well plate. Cells were incubated at 37°C for 7 days, with daily feeding of NIM.

### Lentivirus cloning and production

A lentiviral vector provided by the Brunger lab (Vanderbilt University, Nashville, TN), containing an EFf1α promoter under puromycin resistance, was used as the lentiviral vector backbone. The pCVMV-ACSL1 C-terminally tagged HaloTag plasmid from the Opferman lab (St. Jude Children’s Research Hospital, Memphis, TN) was used as template. Primers for the ACSL1 HaloTag insert (Molecular Cell Biology Resource Core’s Integrated DNA Technologies portal): Forward (5’- GTAATAGCGGCCGCATGCAAGCCCATGAGCTGT-3’) and Reverse (5’-GGCAGCGCTTAATTAAGCCGGAAATCTCGAGCGTC-3’). Primers were designed with overhangs including NotI and PacI restriction cut sites found in the multiple cloning site (MCS) of the lentiviral vector. A polymerase chain reaction (PCR) of 35 cycles was performed on the ACSL1 HaloTag plasmid using the primers above and the Q5 Hot Start High-Fidelity 2X Master Mix (New England Biolabs, cat # M0494S). The PCR cycle included a 95°C denaturation step, a 64°C annealing step, and a 72°C extension step at 15s/kb. The PCR was followed by a double restriction digest of NotI (New England Biolabs, cat # R3189L) and PacI (New England Biolabs, cat # R0547L) on the PCR product of the ACSL1 HaloTag plasmid and the lentiviral vector backbone at 37°C for one hour. The digest was purified via gel extraction (Takara Bio, cat # 740609.250), quantified via Nanodrop, and a 1:3 vector-to-insert molar ratio was used for the ligation step. The digested ACSL1 HaloTag insert and lentiviral backbone were incubated with T4 DNA Ligase (New England Biolabs, cat # M0202L) overnight at 16 °C. The next day, the ligation reaction was treated with Exonuclease V (New England Biolabs, cat # M0345S) for 30 minutes at 37 °C. The treated ligation product was used for transformation of Mix & Go! DH5 Alpha Competent Cells (Zymo Research, cat # T3007). Bacteria were plated on 100 µg/mL ampicillin Luria-Bertani (LB) agar plates, incubated overnight at 37 °C. The following day, bacterial colonies were picked and grown in 10 mL of LB broth overnight at 37 °C, followed by a QIAprep Spin Miniprep (Qiagen, cat # 27104) to isolate plasmid DNA. Sequencing of the clones was performed by GeneWiz, and the corrected lentiviral clones with the ACSL1 HaloTag insert were selected and expanded using the QIAGEN Plasmid Midi Kit (Qiagen, cat # 12143).

### Lentiviral production

A six-well plate was seeded with 700,000 HEK293T cells (Clontech) per well by plating 350,000 cells/mL after dissociating with TrypLE™ Express Enzyme (ThermoFisher Scientific, cat # 12604013). Upon HEK293T cells reaching ∼70-80% confluent, Lipofectamine 3000 (ThermoFisher Scientific, cat # L3000015) was used to transfect 2.0 µg of EF1α ACSL1-HaloTag plasmid, 1.5 µg of pCMVdR8.91 gag/pol packaging plasmid, and 0.6 µg of pMD2.G envelope plasmid (Addgene cat # 12259), following the manufacturer’s instructions. The DNA lipid complex was added dropwise to HEK293T cells cultured with DMEM/F-12 media (Thermofisher Scientific, cat # 11-330-057) and fresh 10% heat- inactivated FBS (Sigma Aldrich, cat # F2442-500ML). One day after transfection, the medium was replaced with 2 mL of Opti-MEM. On days 2 and 3 following transfection, viral medium was collected and filtered with a 0.45 μm PES filter (EMD Millipore, cat # SLFH05010). The supernatant was combined with one volume of Lenti-X Concentrator with 3 volumes of clarified supernatant, mixing by gentle inversion and incubating overnight at 4°C. The sample was centrifuged at 1,500 x g for 45 minutes at 4°C. The pellet was resuspended in 550 uL of Opti- MEM media, and aliquots were stored at −80°C for long-term storage.

### Transduction

For transduction of hESCs, one confluent 35 mm well was dissociated by incubation with 1 mL of Gentle Cell Dissociation Reagent (Stem Cell Technologies, cat # 100- 0485) for 8 minutes at 37°C. Dissociation was quenched with 2 mL of DMEM/F-12 supplemented with 10 µM ROCKi (Stem Cell Technologies cat #72307), and cells were collected into a 15 mL conical tube. Cells were pelleted by centrifugation at 200 × g for 4 minutes, the supernatant was aspirated, and the pellet was resuspended in 1 mL of mTeSR1 (Stem Cell Technologies, cat # 85857) supplemented with 10 µM ROCKi. Lentivirus was added at a final concentration of 5% (v/v) with 4 ug/mL of polybrene (Sigma Aldrich, cat # H9268-5G), cells were plated onto a 24-well plate and incubated overnight at 37°C (16–18 hours).

The following morning, wells were replenished with fresh mTeSR1 and incubated overnight at 37°C. After 48 hours, antibiotic selection began with puromycin at 10 µg/mL (Stem Cell Technologies, cat # 73342). Media was replaced daily with puromycin-containing mTeSR1, and cells were passaged as needed to maintain selection.

### Immunoblot analysis

Cells were harvested in lysis buffer containing 1% Triton X-100 (Sigma, cat # T9284), 1X complete protease inhibitor cocktail (Roche, cat # 04693159001), 1X PhosSTOP phosphatase inhibitor (Roche, cat # 04906837001), and 0.5mM PMSF (RPI, cat # P20270). Samples were kept on ice and vortexed for 30 seconds every 10 minutes. Lysates were centrifuged at 14,100 x g for 30 minutes at 4°C. Supernatant was collected and protein concentrations were quantified using the Pierce BCA Protein Assay Kit (ThermoScientific, cat # 23225). Western blotting was performed using the Bio-Rad Mini-PROTEAN Electrophoresis System (Bio-Rad, cat # 1658036). Samples were run on 4-20% Mini-PROTEAN TGX gels (Bio-Rad, cat # 4561095). Protein was transferred from gels onto methanol activated PVDF membranes (Bio-Rad, cat # 1620177). All membranes were washed in Tris-buffered saline with 0.1% Tween-20 (TBST) and blocked in TBST containing 5% milk for 1 hour at room temperature. Primary antibodies were incubated at listed dilution in TBST containing 5% milk overnight at 4°C. Secondary antibodies were used at listed dilution in TBST containing 5% milk and incubated for 1 hour at room temperature. Protein signal was developed using either Pierce ECL Western Blot Substrate (32109, ThermoScientific) or SuperSignal West Femto (34094, ThermoScientific), depending on the sensitivity of the primary antibody. Protein signal was visualized using the Amersham Imager 600 (29-0834-61, GE Life Sciences). Acquired images were quantified using Thermo Fisher Scientific iBright Analysis Software and representative images were composed using Affinity Designer 2.

| Target | Antibody Name | Host species | Vendor | Catalog Number | RRID | Application |
| --- | --- | --- | --- | --- | --- | --- |
| **ACSL1** | ACSL1 (D2H5) | rabbit | Cell Signaling Technology | 9189S | AB_10891616 | Western blot |
| **beta 3 Tubulin** | Beta III Tubulin | rabbit | Abcam | ab18207 | AB_444319 | IF |
| **beta Actin** | anti-b-Actin | mouse | Sigma-Aldrich | A1978 | AB_476692 | Western blot |
| **CPT1a** | anti-CPT1a | mouse | Proteintech | 66039-01-Ig | AB_2920275 | Western blot |
| **CPT2** | anti-CPT2 | rabbit | Abcam | ab181114 | AB_2687503 | Western blot |
| **MCL-1** | anti-MCL-1 | rabbit | Cell Signaling Technology | 94296S | AB_2722740 | Western blot |
| **MIC10** | MINOS1 (OTI4H7) | mouse | NovusBio | NBP2-45550 | AB_3309700 | Western blot |
| **MIC13** | anti-C19orf70 | rabbit | Proteintech | 25514-1-AP | AB_2880112 | Western blot |
| **MIC19** | anti-CHCHD3 | rabbit | Proteintech | 25625-1-AP | AB_2687533 | Western blot |
| **MIC25** | anti-CHCHD6 | rabbit | Proteintech | 20639-1-AP | AB_10697667 | Western blot |
| **MIC60** | anti-IMMT | rabbit | Atlas Antibodies | HPA036164 | AB_2674979 | Western blot |
| **Mitochondria** | anti-mitochondria | mouse | Abcam | ab92824 | AB_10562769 | IF |
| **OPA1** | anti-OPA-1 | rabbit | Cell Signaling Technology | 67589S | AB_2799728 | Western blot |
| **PAX6** | anti-PAX6 | mouse | BD Biosciences | 561462 | AB_10715442 | IF |
| **PAX6** | PAX6 (D3A9V) XP | rabbit | Cell Signaling Technology | 60433S | AB_2797599 | Western blot |
| **SOX2** | SOX2 (D6D9) XP | rabbit | Cell Signaling Technology | 3579S | AB_2195767 | Western blot |
| **TBR2** | anti-TBR2 | rabbit | Millipore | AB2293 | AB_10806889 | IF |
| **VLCAD** | ACADVL | mouse | Thermo Fisher | PA5-29959 | AB_2547433 | Western blot |

### RNA extraction and cDNA synthesis

Cells were collected in 1000 μl TRIzol reagent. 200 μl of chloroform was added and the samples were incubated at room temperature for 5 minutes prior to centrifugation at 12,000 x g at 4°C. The aqueous phase was collected and 500 μl of isopropanol was added to precipitate RNA. Samples were incubated for 25 minutes at room temperature, followed by centrifugation at 12,000 x g for 10 minutes at 4°C. The RNA pellet was washed with 75% ethanol, semi-dried, and resuspended in 30 μl of DEPC-treated water. Concentration was measured and 2 μg per sample was treated with DNAse (New England Biolabs #M0303) prior to generating cDNA using the manufacturer’s protocol (Thermo Fisher #4368814).

### Real-time quantitative polymerase chain reaction (RT-qPCR)

50 ng of cDNA sample was used to run RT-qPCR using 20 µM of the primers listed in the table. QuantStudio 3 Real-Time PCR machine, SYBR green master mix (Thermo Fisher #4364346), and manufacturer instructions were used to set up the assay.

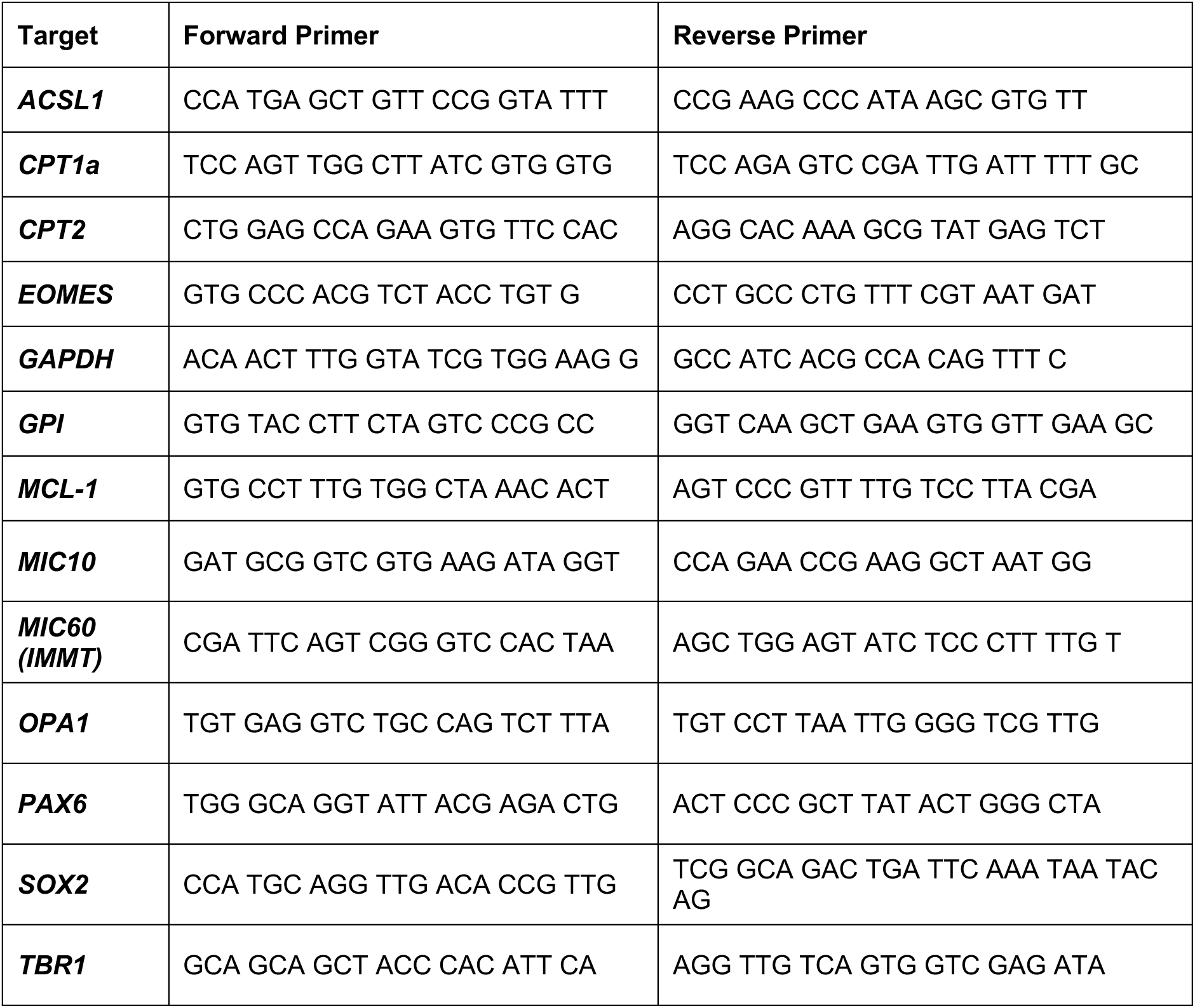
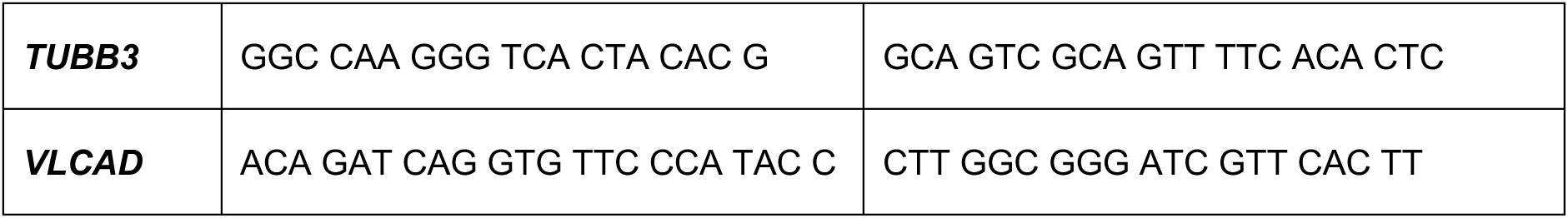

### Immunostaining

Cells were plated on 35 mm glass-bottom dishes (glass thickness of 0.16-0.19 mm) for confocal spinning disk immunofluorescence. Cells were fixed with 4% paraformaldehyde for 20 min and permeabilized in 1% Triton-X-100 for 10 min at room temperature. After blocking in 10% BSA for 1 hour, cells were treated with primary and secondary antibodies using standard methods. Primary antibodies were incubated at listed dilution in 10% BSA overnight at 4°C. Secondary antibodies were used at listed dilution in 10% BSA and incubated for 1 hour at room temperature. The nuclear stain used was HOECHST (Thermo Scientific, cat # H3570) and cells were mounted in Fluoromount-G (Electron Microscopy Sciences #17984- 25) prior to imaging. Cells were labeled for EdU using the manufacturer’s protocol (Thermo Fisher #C10340). Lipid droplet staining was done live using BODIPY (Cayman Chemical Company #25892). Cells were incubated at 37°C for 15 minutes in 4 µM BODIPY prior to fixation in 4% PFA.

### Spinning disk confocal image acquisition

Images were acquired using a Nikon Spinning Disk Confocal outfitted with a Yokogawa CSU- X1 spinning disk head, Andor DU-897 EMCCD camera, and high-speed piezo [z] stage. All 100X images were acquired with a step size of 0.2 μm, while 20X images were acquired with a step size of 2 µm. Image processing and quantification was performed using NIS Elements (Nikon) and all representative images were generated using Fiji.

Quantification of mitochondrial morphology was performed in NIS-Elements version 5.42.06 (Nikon). Mitochondria were segmented in 3D, and data were extracted from the resulting 3D mask. Major axis length, surface area, volume, and count measurements were exported into Excel and data was analyzed using GraphPad Prism v9.

### Tissue preparation for transmission electron microscopy (TEM) and focused ion beam scanning electron microscopy (FIB-SEM)

Neural progenitor cells were plated at 1.5x10^6^ cells per 35 mm dish on Matrigel. The cells were washed with 1X PBS (3X), fixed in 2.5% glutaraldehyde in 0.1 M cacodylate prewarmed to 37°C, left to cool down at room temperature for 1hr, and placed at 4°C for an additional 24-48 hour of fixation. Cells were washed from fixative with 1X PBS (3x). Samples were sequentially post-fixed using the OTO method: 1.5% potassium ferrocyanide, 1% OsO4 in 0.1 M cacodylate for 30 minutes on ice, then 1% thiocarbohydrazide for 10 minutes, followed by 1% OsO4 in ddH2O for 10 minutes. Samples are then in-block stained with 2% uranyl acetate followed by lead aspartate. The samples were then dehydrated in a graded ethanol series and infiltrated with Durcupan resin using acetone as the transition solvent. The resin was polymerized at 60°C for 48 hours. For TEM, sections were cut on a Leica UC7 at a nominal thickness of 50 nm with no post-section staining. For FIB-SEM, samples were prepared as previously described^63^.

### TEM image acquisition and cristae scoring

Sections were imaged with either a ThermoFisher Scientific Tecnai T12 operating at 100 keV using an AMT nanosprint5 CMOS camera or a JEOL 2100+ operating at 200 keV using an AMT nanosprint15 II CMOS camera. In both cases, AMT acquisition software was used for imaging. The TEM images were analyzed using cristae scoring^64^. The files were first renamed to blind the scoring process. The images were then blindly scored. A score was provided for each mitochondrion based on the quality of the cristae. The scoring rubric was as follows: 0 - there were no crista, 1 - more than 50% of the mitochondrial area did not have cristae, 2 - more than 25% of the mitochondria was without cristae, 3 - lots of cristae but irregular in structure, and 4 - many cristae that are regular structures.

### FIB-SEM image acquisition

Sections were imaged with Zeiss Crossbeam 550 FIB-SEM using ATLAS5 software for automated volumetric acquisition. Sample blocked were polished with a diamond knife and coated with platinum. Platinum pads were deposited via GIS on ROIs of interest, followed by milling in tracking marks and coating with GIS carbon. Milling was performed using a 700 pA Ga beam at 8 nm slice depth, imaging was performed 1.5 keV electron beam using the in column ESB detector for imaging at 8 nm pixels to make isotropic voxels. Volumetric slice alignments were performed in ATLAS5 using the two-window alignment method.

### FIB-SEM mitochondrial image analysis

FIB-SEM stacks were first cropped using Fiji is just Image J (FIJI). In the FIB-SEM stacks, none of the cells imaged were able to be visualized completely. Therefore, the crop size and location were selected by choosing a cell with bright, visible mitochondria. The crop was 500 x 500 pixels with a 3,000 nm depth. The cropped image was then imported into the FIJI LabKit plugin for segmentation^65^. For segmentation, each mitochondrion was manually traced with a separate label ID. Quality control was then performed following each mitochondrion through the entirety of the Z-stack and tracking all locations where the membranes connect. Any mitochondria that merged in any of the frames were labeled with the same label ID. The cropped FIB-SEM image and the proofread segmentation masks were imported into the IMARIS software. Ortho slices in the XY and XZ planes were added, and the proofread mask was selected to be identified by ID. This identifies each separate mitochondrion. Any mitochondria that were still split were manually combined in the IMARIS software using the unify feature. A clipping plane was then added. Animations were made using the animations tab in IMARIS with 50 frames per second and 1,000 frames total. Still images were captured using the screenshot feature. Volume quantification was done by volume measurement provided in the statistics tab in IMARIS. Surface area quantification was manually calculating using volume and sphericity measurements from the IMARIS statistics tab using the equation: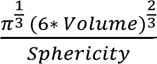. Lastly, the mitochondrial branching index (MBI), was calculated by obtaining the BoundingBox00 length A (longest side) and BoundingBox00 length C (Shortest side) in the Vantage 2D tab and then dividing Length C/ Length A manually^66^.

### FIB-SEM cristae image analysis

The same FIB-SEM cropped images were used for the segmentation of the cristae. The cropped images were imported into FIJI and applied Contrast Limited Adaptive Histogram Equalization (CLAHE) to increase contrast. They were then imported into the LabKit plugin for segmentation. Each crista was manually traced, and all the cristae within one mitochondrion were labeled with the same label ID. Quality control was then performed, ensuring each mitochondrion had all its cristae labeled with the same ID. The cropped FIB-SEM image and the proofread segmentation masks were imported into the IMARIS software. Ortho slices in the XY and XZ planes were added, and the proofread mask was selected by ID. This identifies each separate mitochondrion’s cristae. Any errors in the cristae mask were merged using the unify feature in IMARIS. A clipping plane was then added to show the mitochondria throughout the entire z-plane. Animations were made using the animations tab in IMARIS with 50 frames per second and with 1,000 frames total. Still images were taken using the snapshot feature. Volume quantification was done by volume measurement provided in the statistics tab in IMARIS. Surface Area quantification was manually calculated using volume and sphericity measurements from IMARIS statistics tab using the equation: 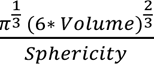. Cristae measurements were then normalized to the corresponding mitochondria by dividing the crista measurement by the mitochondrion measurement 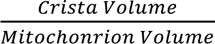 . The mitochondria segmentation was then added to the same IMARIS file as a second surface. The label was chosen by ID, and the surface type was changed to transparent to show the cristae within those mitochondria. Images were taken using the snapshot function, and animations were made with 50 frames per second and 1,000 frames total.

### Statistical analysis

All experiments were performed with at least 3 biological replicates. Statistical significance was determined by one-way, repeated measures one-way ANOVA, or t-test as appropriate for each experiment. Significance was assessed using Fisher’s protected least significance difference test.

GraphPad Prism v9 was used for most statistical analyses and data visualization, unless otherwise specified in the figure legend. Error bars in all bar graphs represent the standard error of the mean unless otherwise noted in the figure. No outliers were removed from the analyses. For all statistical analyses, a significant difference was accepted when P < 0.05.

### Modeling *Ct* from *RT-qPCR* data

Cycles to threshold, 𝐶𝑡, measured using *RT-qPCR* can be modeled as previously shown^67^,

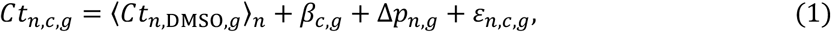

where 𝑛 is the biological replicate, 𝑐 is the treatment condition, and 𝑔 is the gene of interest. The first term, 〈𝐶𝑡_𝑛,DMSO,g_〉_𝑛_, is 𝐶𝑡 for gene 𝑔 under control condition DMSO averaged across all biological replicates. The next term, coefficient 𝛽_𝑐,g_, quantifies the effect of treatment 𝑐 on gene 𝑔. Coefficient Δ𝑝_𝑛,g_ = 𝑝_𝑛,g_ − 𝑝_1,g_ is a plate effect, allowing for differences between plates 𝑛 for gene *g*^68^. To maintain parameter identifiability, the plate effect is defined as the difference between plate 𝑛 and plate 𝑛 = 1. Plate number is indicated by the same index as the biological replicates since each plate contains a single biological replicate. The remaining term 𝜀_𝑛,𝑐,g_models unobserved random errors.

From this we derive the quantity of interest, ΔΔ𝐶𝑡_𝑛,𝑐,g_, given by,

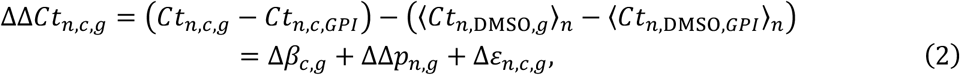

where Δ𝛽_𝑐,g_ = 𝛽_𝑐,g_ − 𝛽_𝑐,𝐺𝑃𝐼_, ΔΔ𝑝_𝑛,g_ = Δ𝑝_𝑛,g_ − Δ𝑝_𝑛,𝐺𝑃𝐼_, and Δ𝜀_𝑛,𝑐,g_ = 𝜀_𝑛,𝑐,g_ − 𝜀_𝑛,𝑐,𝐺𝑃𝐼_. We fit this model to measured data using weighted linear regression with the Python package stats models [available at: https://proceedings.scipy.org/articles/Majora-92bf1922-011]. We assume that intra-gene variance is approximately homoscedastic, while inter-gene variance is heteroscedastic. Thus, weights are set at the gene level, giving,

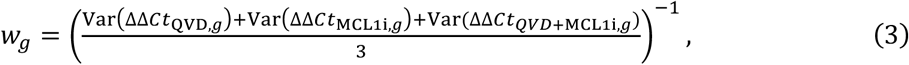

where 𝑉𝑎𝑟() is the sample variance for gene 𝑔 and indicated treatment across all biological replicates.

## Acknowledgements

We thank Caleb Hayes, Caroline Bodnya, Meeros Khattaie, Dr. Antentor J. Hinton, Dr. Zer Vue, Dr. Evan Krystofiak and Dr. Oleg Kovtun for their technical advice and assistance. This work was supported by 2R35 GM128915-01NIGMS (VG), Training Program in Stem Cell and Regenerative Developmental Biology 5T32HD007502-25 (MRH), and HHMI Gilliam Fellowship GT15720 (MG). All TEM and FIB-SEM imaging and image analysis were performed in part using the Vanderbilt Cell Imaging Shared Resource, which is supported by NIH grants 1S10OD012324-01 and 1S10OD021630-01. The authors declare no competing financial interests.

## Author contributions

M. R. Hanna and V. Gama conceived the study, designed experiments, interpreted data, and wrote the manuscript. V. Gama supervised the research. M. R. Hanna designed, conducted, and analyzed all cell biology experiments, with technical assistance from M. Gil, M. Yarbrough, and J. Costanzo. M. Yarbrough performed 3D reconstruction and analysis of FIB-SEM data. M. Murrow performed statistical analysis and modeling for some of the data. All authors edited the document.

