## Supplemental Figures and Legends for "Pharmacological MCL-1 inhibition disrupts fatty acid oxidation and depletes neural progenitor cells"

**Supplementary Figure Legends**

**Supplementary Figure 1. S63845 effectively inhibits MCL-1 in human NPCs.** (**A**) Quantified relative expression measured by RT-qPCR of *MCL-1* after 24- and 48-hour treatment with S63845 (MCL-1 inhibitor) -/+ QVD (caspase inhibitor). Two technical replicates were used for all RT-qPCR data. RT-qPCR data are displayed as relative fold expression derived from $2^{(-\Delta\Delta CT)}$, and raw data were analyzed using ANCOVA as described in the methods section. (**B**) Representative images (n=3) (**C**) Quantified band density normalized to Actin loading control of MCL-1 after 24- and 48-hour treatments. Each data point on the graphs represents a biological replicate. Band density data were analyzed using an ordinary two-way ANOVA with an uncorrected Fisher's LSD test for multiple comparisons. Shape of the data points corresponds to a biological replicate (circle for n=1, square for n=2, and triangle for n=3). All tests used a confidence level of 95% (p-value ≤ 0.05 as the significance threshold). Error bars represent standard error of the mean.

**Supplementary Figure 2. S63845 does not alter mitochondria branching index and sphericity.** (**A**) FIB-SEM 3-dimensional reconstruction of mitochondria after NPCs were treated with S63845 (MCL-1 inhibitor) for 48h (scale bar = 3000 nm). (**B**) Mitochondrial branching index (MBI) and sphericity were quantified as described in the methods section. Each data point represents an individual instance of a mitochondrion. Data analyzed using a parametric two-tailed unpaired t-test with Welch's Correction and a 95% confidence level (p-value ≤ 0.05 as the significance threshold). Error bars represent standard error of the mean.

**Supplementary Figure 3. S63845 induces changes subcomplex MICOS proteins.** (**A**) Representative images (n=3) (**B**) Quantified band density normalized to Actin loading control of subcomplex MICOS proteins, MIC19, MIC25, and MIC13, after 24- and 48-hours of treatment with S63845 (MCL-1 inhibitor) -/+ QVD (caspase inhibitor). Each data point represents a biological replicate. Shape of the data points corresponds to a biological replicate (circle for n=1, square for n=2, and triangle for n=3). Data were analyzed using an ordinary two-way ANOVA with an uncorrected Fisher's LSD test for multiple comparisons and a 95% confidence level (p-value ≤ 0.05 as the significance threshold). Error bars represent standard error of the mean.

**Supplementary Figure 4. S63845 impairs mitochondrial respiration in human NPCs at 24 and 48 hours.** (**A**) Oxygen consumption rate (OCR) and extracellular acidification rate (ECAR) traces over time following 24-hour treatment with S63845 (MCL-1 inhibitor) -/+ QVD (caspase inhibitor), measured by Seahorse XF Mitochondrial Stress Assay. (**B**) Quantified metabolic parameters derived from OCR trace for 24h treatments, including basal respiration, maximal respiration, ATP-linked respiration, proton leak, coupling efficiency, non-mitochondrial OCR, spare respiratory capacity, and spare respiratory capacity (%), shown for DMSO and S63845-treated cells -/+ QVD. (**C**) OCR and ECAR traces over time following 48-hour treatment with S63845 -/+ QVD. (**D**) Quantified metabolic parameters derived from OCR trace 48h treatments, including basal respiration, maximal respiration, ATP-linked production, proton leak, coupling efficiency, non-mitochondrial OCR, spare respiratory capacity, and spare respiratory capacity (%), shown for DMSO and S63845-treated cells -/+ QVD. Each data point on the graphs represents a biological replicate. Shape of the data points corresponds to a biological replicate (circle for n=1, square for n=2, and triangle for n=3). Data were analyzed using an ordinary two-way ANOVA with an uncorrected Fisher's LSD test for multiple comparisons and a 95% confidence level (p-value ≤ 0.05 as the significance threshold). Error bars represent standard error of the mean.

**Supplementary Figure 5. Downregulation of key NPC identity markers caused by S63845 persists in the absence of caspase-mediated cell death.** (**A**) Representative images (n=3) and quantified band density normalized to Actin loading control of NPC identity markers, PAX6 and SOX2 after treatment with S63845 (MCL-1 inhibitor) -/+ QVD (caspase inhibitor). Band density data were analyzed using an ordinary two-way ANOVA with an uncorrected Fisher's LSD test for multiple comparisons. (**B**) Quantified relative expression measured by RT-qPCR of identity markers, *SOX2*, *PAX6*, *EOMES*, *TBR1*, and *TUBB3*, after 24- (top panel) and 48-hour (bottom panel) treatments. Two technical replicates were used for all RT-qPCR data. RT-qPCR data are displayed as relative fold expression derived from $2^{(-\Delta\Delta CT)}$, and raw data were analyzed using ANCOVA as described in the methods section. Each data point represents a biological replicate. Shape of the data points corresponds to a biological replicate (circle for n=1, square for n=2, and triangle for n=3). All tests used a confidence level of 95% (p-value ≤ 0.05 as the significance threshold). Error bars represent standard error of the mean.

**Supplementary Figure 6. S63845 does not affect hNPC proliferation.** (**A**) Immunofluorescent images of proliferating cells (EdU in white) and nuclei (Hoechst in cyan) acquired on SDC microscope at 60X (scale bar = 50μm). Representative images (n=3) of hNPCs treated with S63845 (MCL-1 inhibitor) -/+ QVD (caspase inhibitor) after 24- (left panel) and 48-hours (right panel). (**B**) Percent of EdU-labeled hNPCs after 24- (top panel) and 48-hour (bottom panel) treatments. Graphs from left to right: cell count of total proliferating cells, early S-phase (whole nuclear labeling), and late S-phase (punctate nuclear labeling) normalized to total cell count; cell count of early S-phase, and late S-phase normalized to total proliferating cells. Each large, opened-shape data point (black) represents the mean of a biological replicate, and each small, closed-shape data point (transparent gray) represents a technical replicate (a single image). Shape of the data points corresponds to a biological replicate (circle for n=1, square for n=2, and triangle for n=3). Data were analyzed using an ordinary two-way ANOVA with an uncorrected Fisher's LSD test for multiple comparisons and a 95% confidence level (p-value ≤ 0.05 as the significance threshold). Error bars represent standard error of the mean.

#### Supplementary Figure1

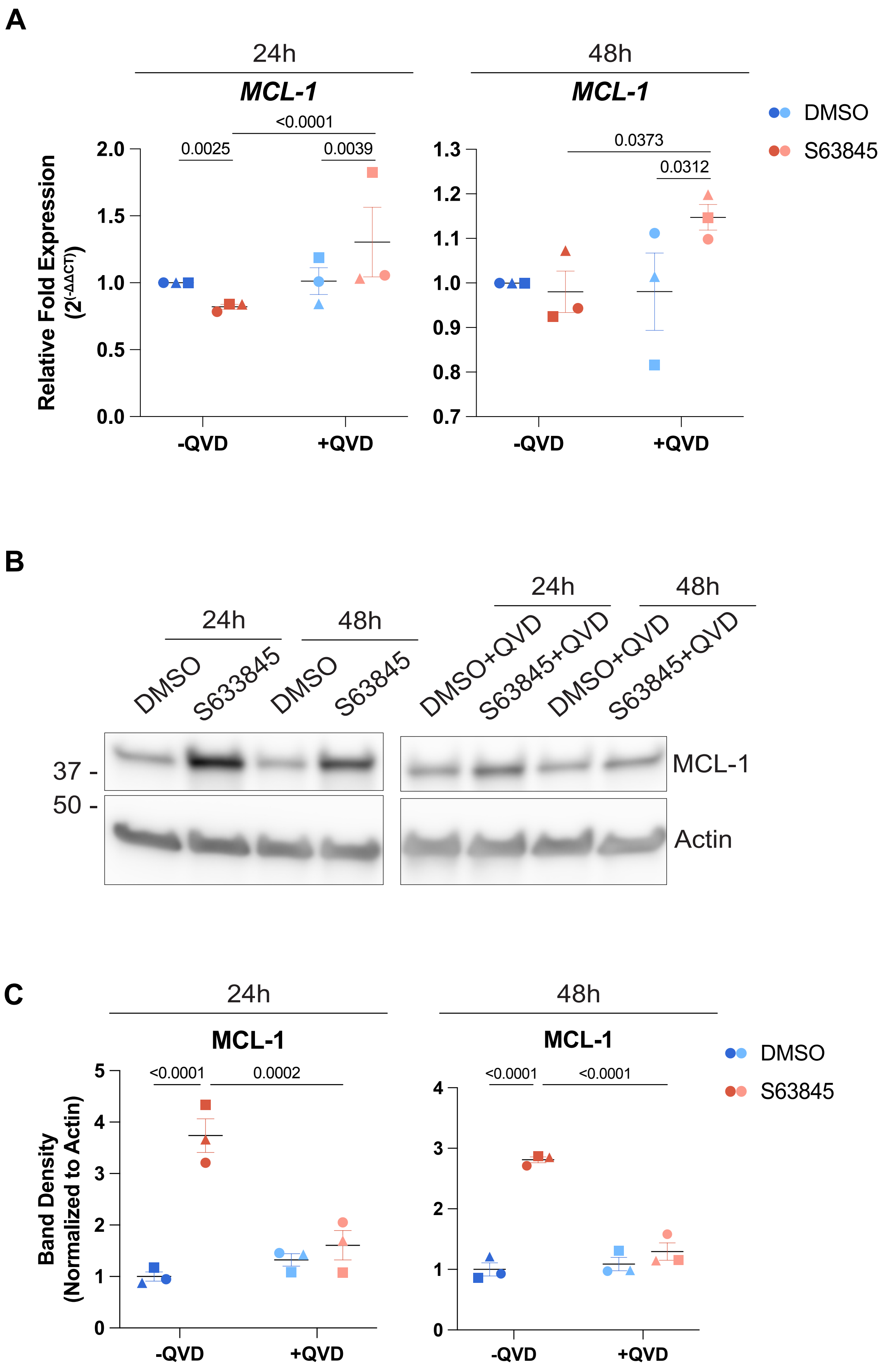

**Supplementary Figure 2**

A

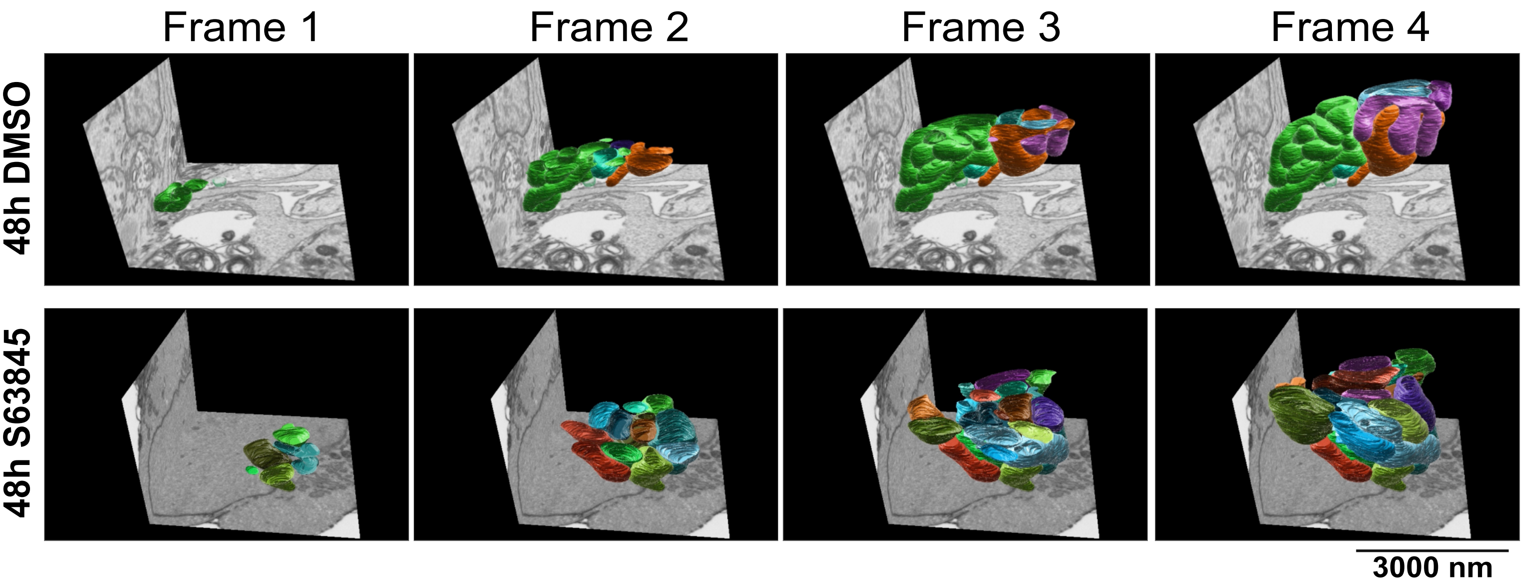


B

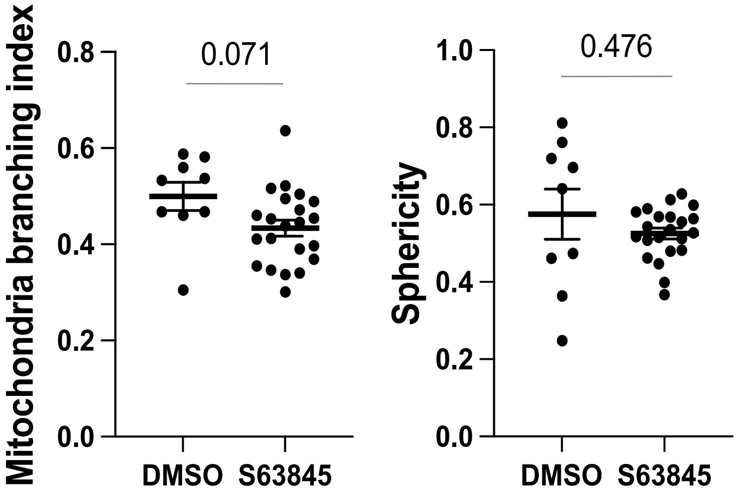

#### Supplementary Figure 3

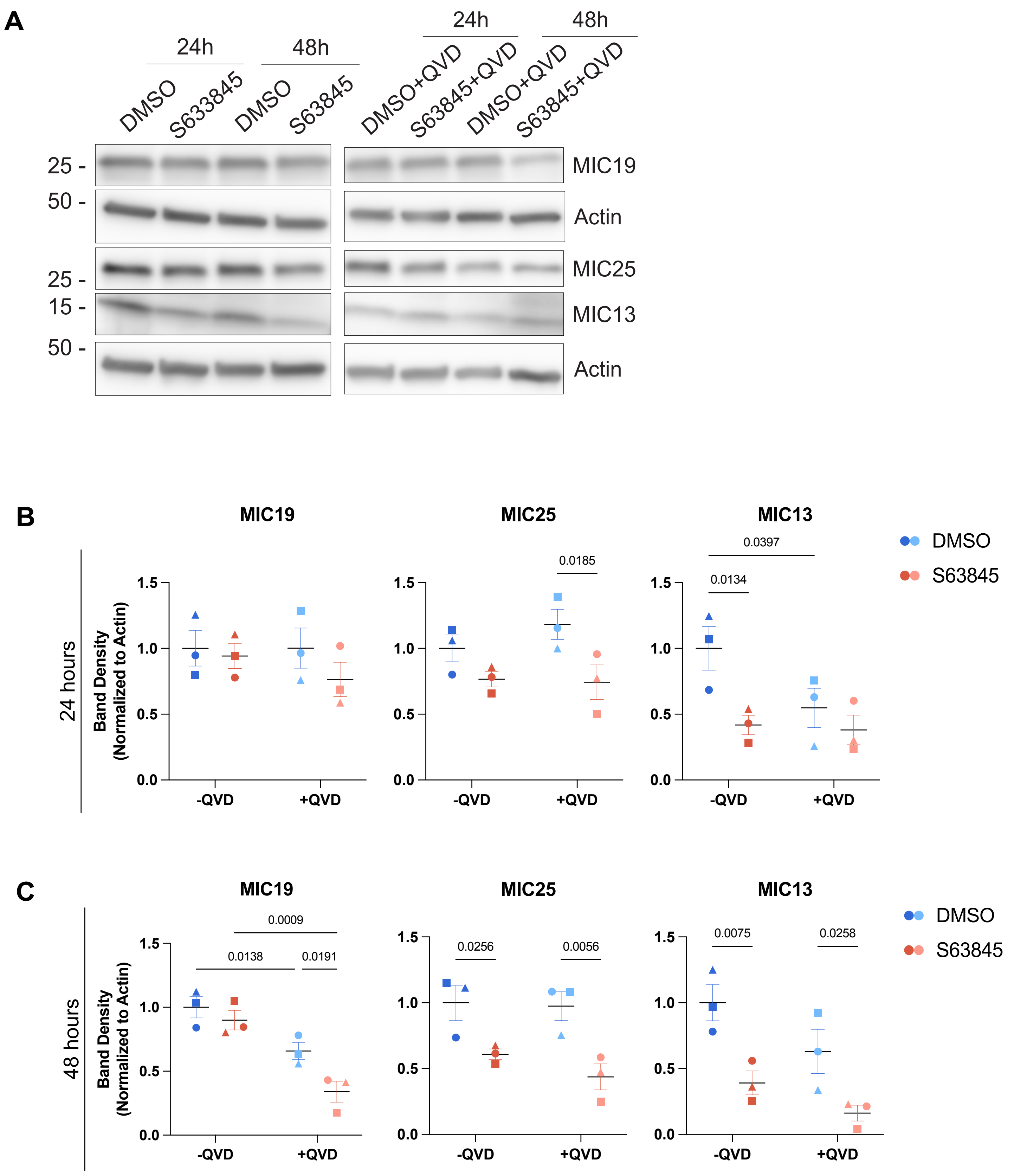

### Supplementary Figure 4

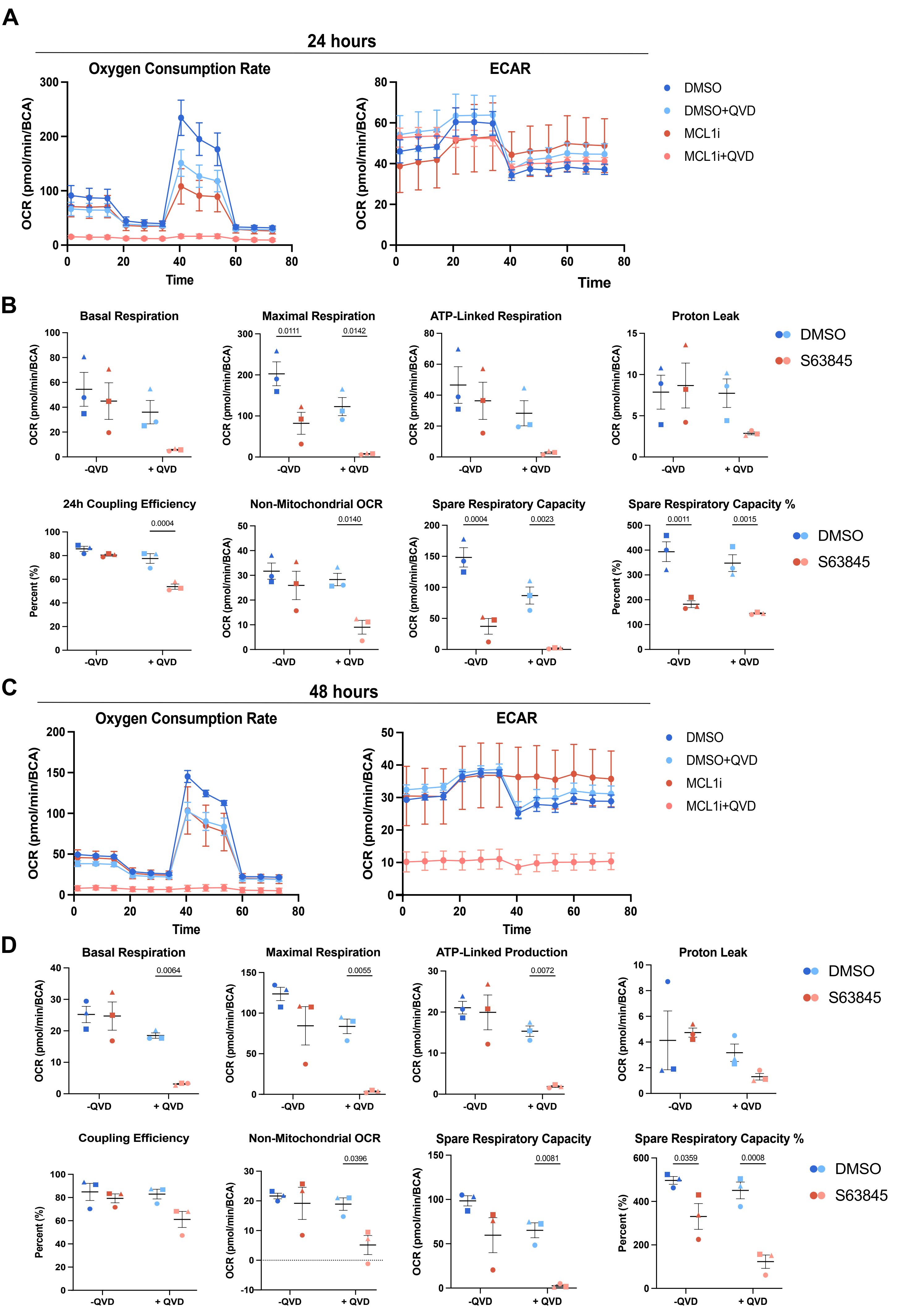

### Supplementary Figure 5

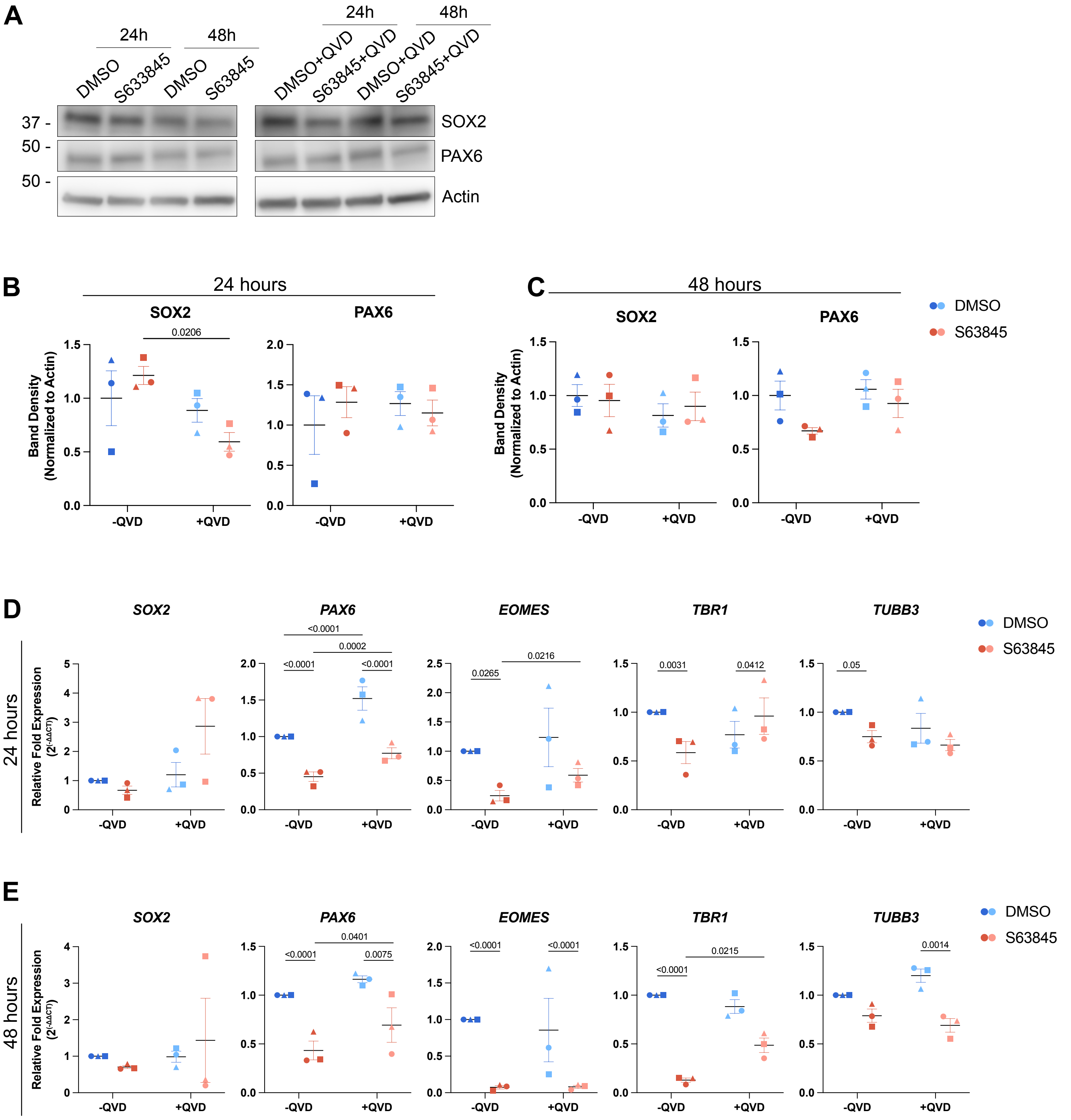

#### Supplementary Figure 6

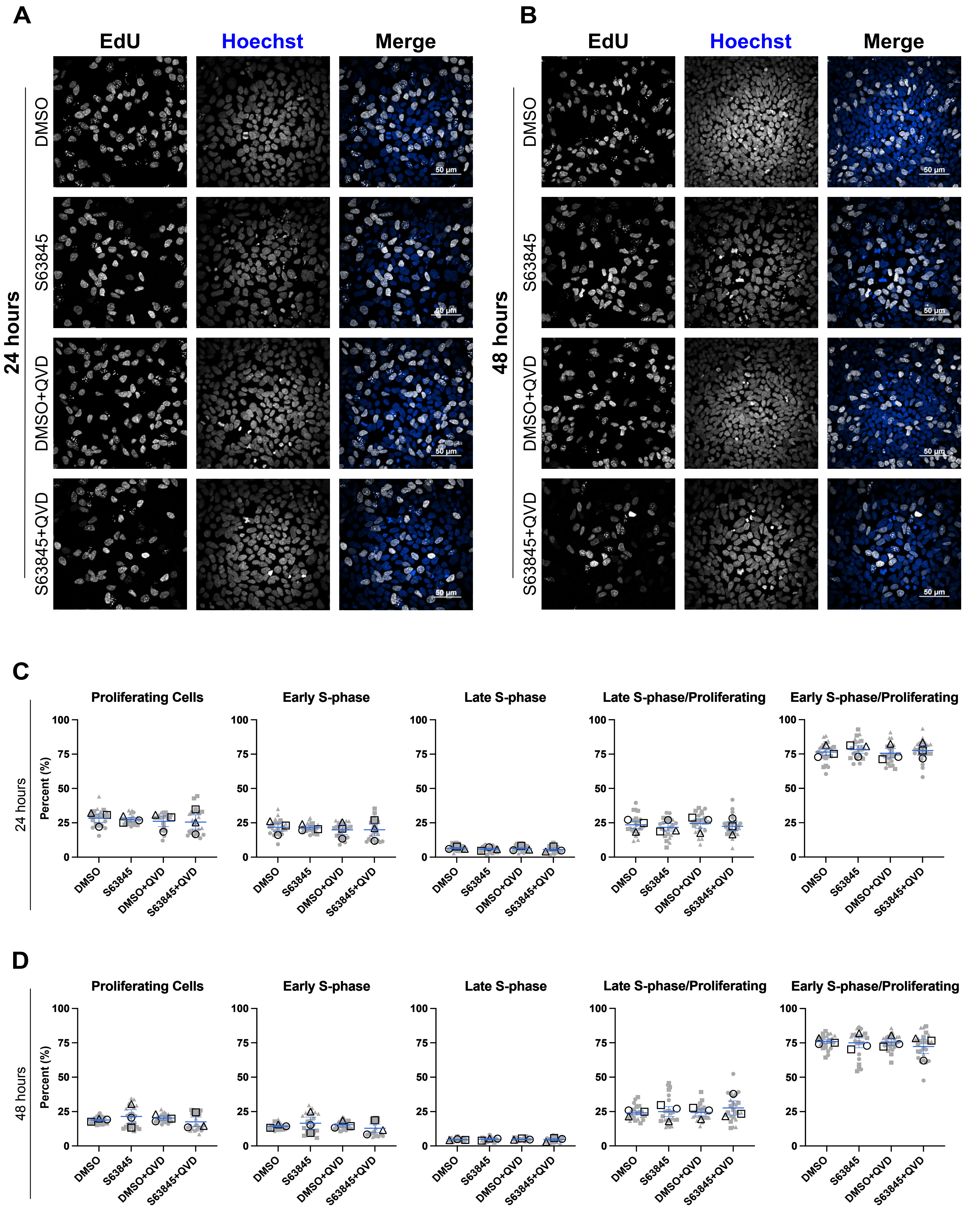
